# Tissue-wide integration of mechanical cues promotes efficient auxin patterning

**DOI:** 10.1101/820837

**Authors:** João R. D. Ramos, Alexis Maizel, Karen Alim

## Abstract

New plants organs form by local accumulation of auxin, which is transported by PIN proteins that localize following mechanical stresses. As auxin itself modifies tissue mechanics, a feedback loop between tissue mechanics and auxin patterning unfolds – yet the impact of tissue-wide mechanical coupling on auxin pattern emergence remains unclear. Here, we use a hybrid model composed of a vertex model for plant tissue mechanics, and a compartment model for auxin transport to explore the collective mechanical response of the tissue to auxin patterns and how it feeds back onto auxin transport. We compare a model accounting for a tissue-wide mechanical integration to a model where mechanical stresses are averaged out across the tissue. We show that only tissue-wide mechanical coupling leads to focused auxin spots, which we show to result from the formation of a circumferential stress field around these spots, self-reinforcing PIN polarity and auxin accumulation.

## Introduction

Formation of organs entails an effective coordination of local cell growth typically initiated by patterns of one or more morphogenic factors. Understanding how these patterns of morphogenic agents robustly emerge is fundamental. Plants organs formation is interesting from a physical perspective due to the strong mechanical coupling between plant cells, and the fact that growth is driven by changes in the mechanical properties of the cell wall and internal pressure (1–5). Evidences indicate that the morphogenic factors changes the mechanics of the tissue (6, 7), with implications for the shaping of organs (8). Interestingly, the transporters of these factors respond to mechanical cues (9, 10), leading to a intertwining of chemical and mechanical cues.

The key morphogenic agent in plants is the phytohormone auxin, Indole-3-Acetic Acid. Auxin accumulation drives a wide range of plant developmental processes including, but not limited to initiation of cell growth, cell division, and cell differentiation (11–13). Establishment of auxin patterns is ubiquitous in plant organ morphogenesis (14). The best characterized example are the regular patterns of auxin spots in the outmost epidermal cell layer at the tip of the shoot that prefigures the regular disposition of organs called phyllotactic pattern (15–19). These auxin accumulation spots mark the location of emerging primordia of new aerial plant organs. Auxin patterns result from the polar distribution of auxin efflux carriers called PIN-FORMED (PINs) (14, 15, 20–22). Because of its prevalence in plant development, understanding how these auxin patterns emerge has been intensively studied and mathematically modelled. Auxin concentration feedback models (23–27), organize their flow up-the-gradient of auxin concentration, reinforcing auxin maxima. Canalization models, or flux-based models, (28–34) reinforce already existing flows, and, as such, both up-the-gradient and down-the-gradient flows can exist. Some attempts at unifying both mechanisms have been made (35–37), yet many conditions have to be imposed to explain, for instance, the fountain-like patterns arising during root development (38).

Tissue mechanics has emerged as a potent regulator of plant development (5, 39–42). Plant cells are able to read mechanical stress and respond accordingly, rearranging their microtubules along the main direction of mechanical stresses (39). Furthermore, PIN1 polarity and microtubules alignment at the shoot apical meristem are correlated (9), suggesting that PINs localisation is regulated by mechanical cues such as strain or stress. This coupling between PINs localisation and mechanical cues is theoretically able to predict PIN polarity and density for a wide range of cell wall stress and membrane tension (10). Such coupling is also supported by several other observations: the physical connection of PINs to the cell wall (43), the change in polarity induced by cell curvature (44) and disorganization of PIN polarity by modification of the cell wall mechanical properties (7).

Auxin can induce remodelling of the cell wall and thus modify its mechanical properties (4, 6, 7, 45). This may in turn influence PINs localisation and therefore have consequences on the pattern of auxin. Modeling of this feedback in a tissue showed that mechanical stresses can lead to the emergence of a regular phyllotactic auxin pattern by regulating PIN localisation (9). Although this result shows the importance of local mechanical coupling (Fig. 1) for emergence of auxin patterns, it remains unknown whether tissue-wide mechanical coupling plays a role in the emergence of these patterns.

**Fig. 1.**
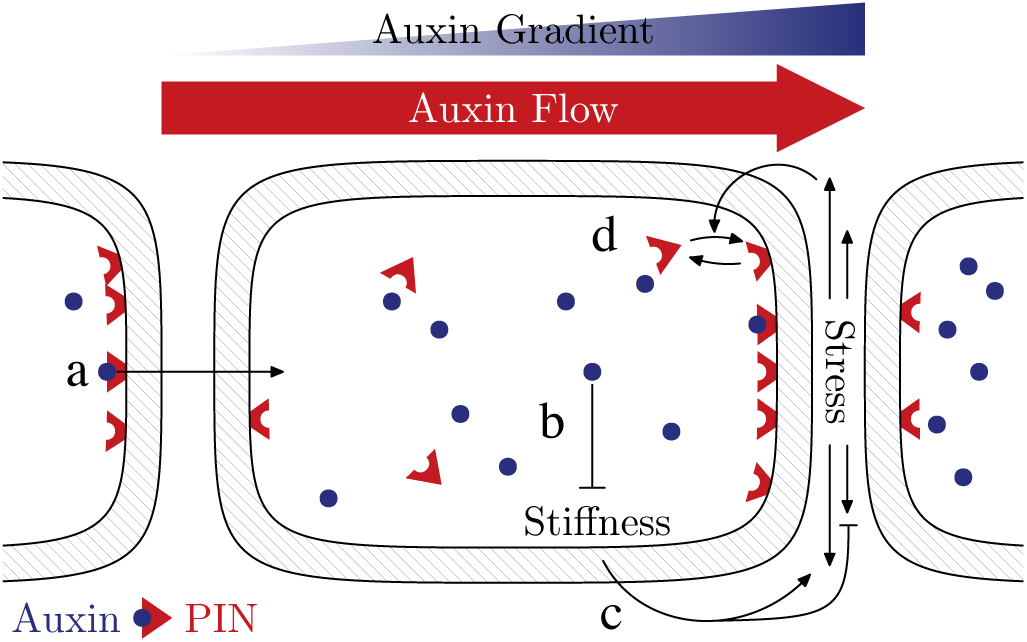
Schematic representation of the cell-cell feedback mechanism between cell wall loosening via auxin and mechanical control of PIN. (a) Auxin is transported to neighbouring cells via bound PIN efflux carriers. (b) Auxin interacts with the mechanical properties of the cell wall reducing its stiffness. (c) Increasing stiffness of a particular wall component shifts the stress load from the component of its neighbour to itself. (d) Wall stress promotes PIN binding. A difference in auxin, therefore, induces a stress difference between the two compartments separating both cells. This stress difference is such that PIN binds preferentially in the cell with lower auxin concentration, increasing the flow of auxin into the cell with higher auxin concentration.

To address if tissue-wide mechanical feedback on auxin patterning promotes or inhibits auxin patterning we adapted the model for auxin transport used in (9) to a vertex model mechanical description of a tissue. We compared a full mechanical description of the tissue, in which each cell is mechanically coupled to all other cells, to a static tissue under constant averaged stress, in which cells are only mechanically coupled only to their direct neighbours through stress load division (Fig. 2). By comparing the patterns in the two models we find that due to stress-fields arising from mechanical feedback the magnitude of auxin spots is larger for lower stress-PIN coupling, indicating that the inclusion of tissue-wide stress patterns is more efficient to reach biologically relevant auxin patterning. Finally, by studying how stress self-organizes according to auxin concentration patterns we find that anisotropic stress fields form around auxin spots reinforcing transport towards auxin maxima.

**Fig. 2.**
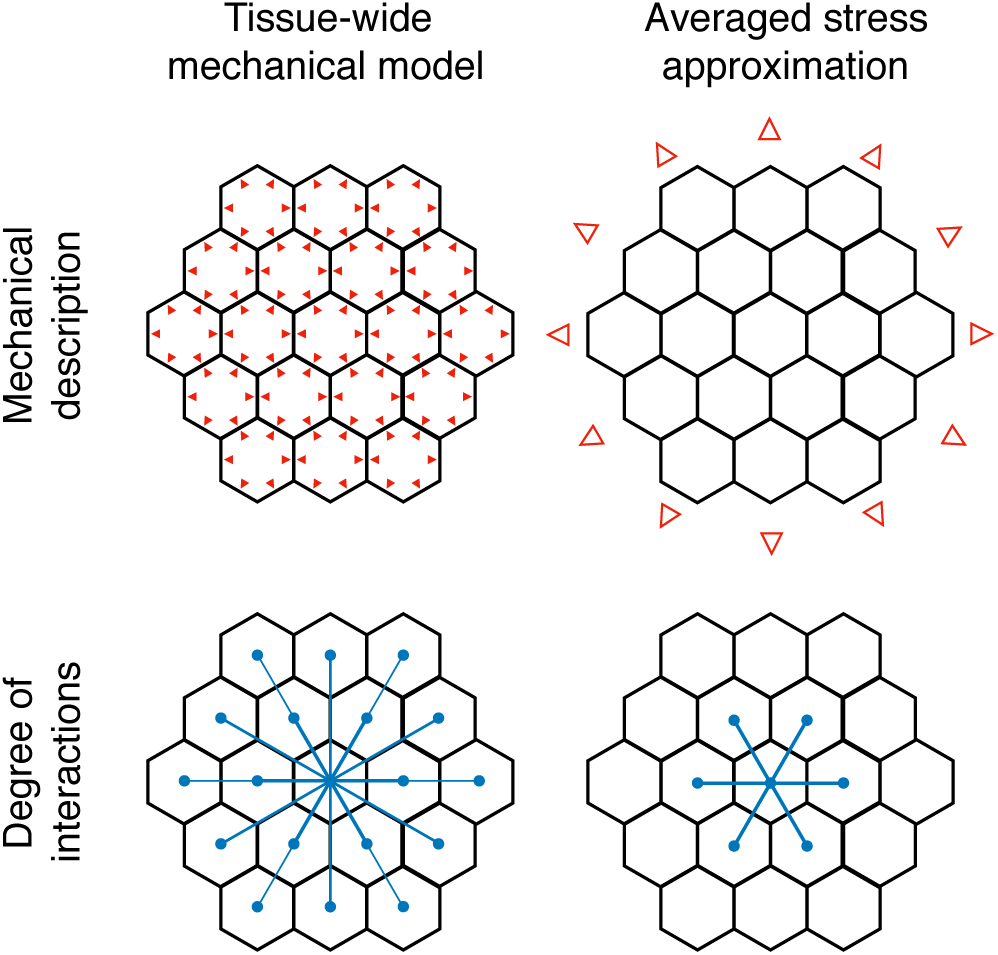
Schematic difference between the two studied regimes. The tissue-wide mechanical model description takes into account the turgor pressure of individual cells, whereas the averaged stress approximation assumes an uniform isotropic stress acting on the whole tissue, resulting in different degrees of interaction. In the first case, wall stress is a function of all other vertices in the tissue (hence the name), whereas in the second case, it is only influenced by the immediate neighbours. Comparison of both models reveal the impact of tissue-wide stress patterns.

## Methods

In order to investigate the interaction between auxin cell wall softening and collective tissue mechanics, we use a vertex model to describe the mechanical behaviour of the tissue, and a compartment model to express auxin concentration and transport between adjacent cells.

### Geometrical setup of the tissue

The tissue is described by a tiling of two dimensional space into *M* cells surrounded by their cell walls. Walls are represented as edges connecting two vertices each, positioned at **x**_*i*_ = (*x*_*i*_, *y*_*i*_), *i*∈ [1, *N*]. Here, we reserve Latin indices for vertex numbering and Greek ones for cells. Each cell wall segment has two compartments, one facing each cell. Therefore, we represent each cell wall with two edges of opposite direction, one for each compartment. The position of tissue vertices fully define geometrical quantities such as cell areas, *A*_*α*_, cell perimeters, *L*_*α*_, wall lengths, *l*_*ij*_ = *l*_*ji*_ and cell centroids, **X**_*α*_ (Fig. 3A). To simplify notation significantly we also define for each cell the cyclically ordered set of all vertices around that cell, 𝒱_*α*_, arranged counterclockwise. Similarly, we introduce 𝒩_*α*_ as the cyclically ordered (counterclockwise) set of all neighboring regions around cell *α*, one for each edge of *α* (Fig. 3B).

**Fig. 3.**
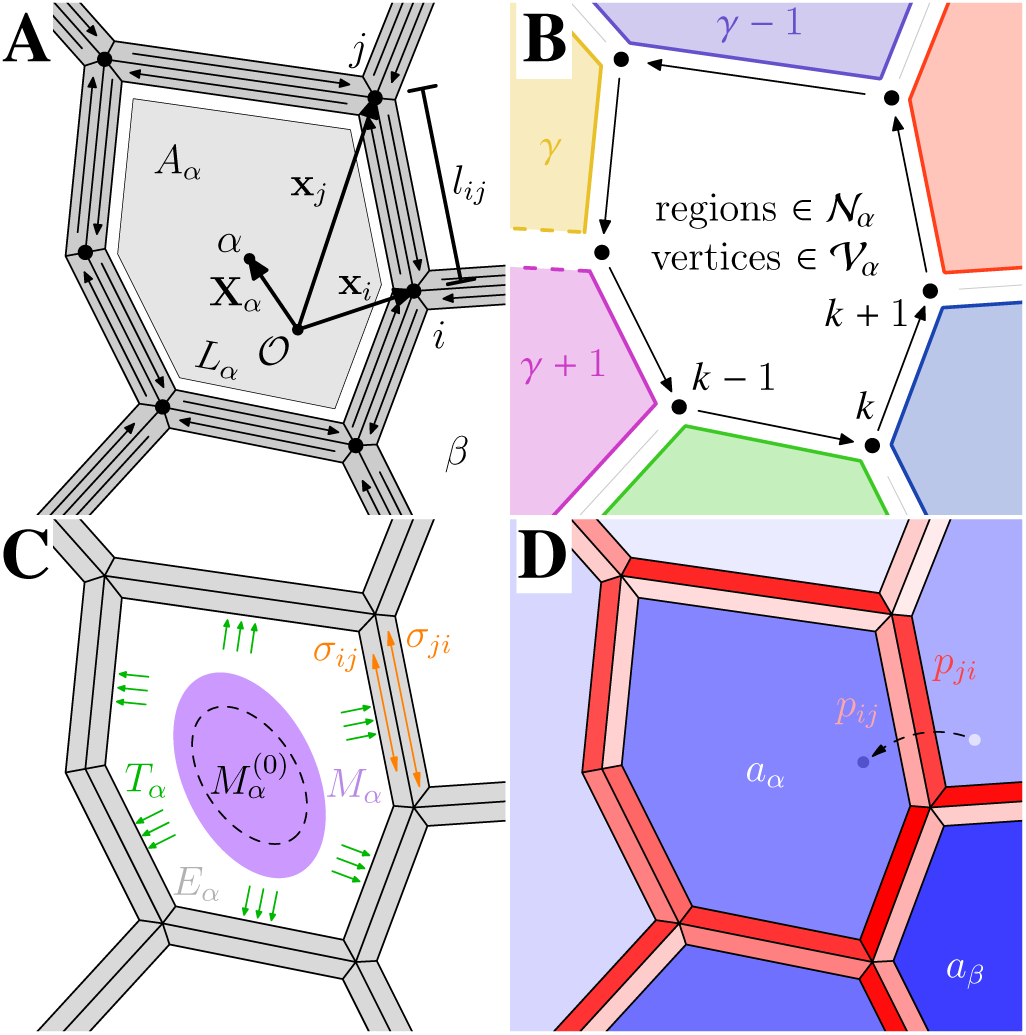
Vertex model description of a cell as a geometrical, mechanical, and biologically active entity. (A) A cell *α* surrounded by its cell walls with centroid **X**_*α*_, area *A*_*α*_ and perimeter *L*_*α*_. Each cell wall has two compartments, one for each adjacent cell. Vertices *i* and *j* have positions **x**_*i*_ and **x** _*j*_ and the distance between them is *l*_*ij*_ = *l*_*ji*_. (B) Set of surrounding regions, one for each wall, 𝒩_*α*_, and set surrounding vertices, 𝒱_*α*_, used in the equations of the model. (C) Mechanically, cell *α* is under turgor pressure *T*_*α*_, the surrounding wall compartments have stiffness *E*_*α*_. *M*_*α*_ is the second moment of area of cell *α*, whereas 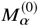 is that same quantity when the cell is at rest. *σ*_*ij*_ refers to the longitudinal stress acting on the compartment of the wall on the immediate left of the directed edge that connects *I* to *j*. (D) Cell *α* has an auxin concentration *a*_*α*_ which is expressed, degraded and transported, both passively and actively. The active component of auxin transport relies on the density of membrane bound efflux auxin carriers facing a particular wall compartment, which we represent by *p*_*ij*_.

### Tissue Mechanics – Tissue-wide Coupling

Vertex models are a widely employed theoretical approach to describe mechanics of epithelial tissues and morphogenesis (46–51). The essence of vertex models is that cell geometry within a tissue is given as the mechanical equilibrium of the tissue. In the case of plant cells, the shape of a cell is a competition between the turgor pressure, *T*_*α*_, all cells exert on each other and the cell’s resistance to deformation with stiffness, *E*_*α*_. Strain acting on each cell will be described using the second moment of area of the corresponding cell in reference to its centroid, *M*_*α*_, whose components are

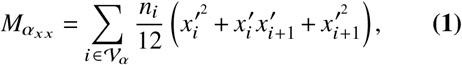

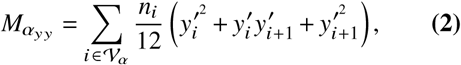

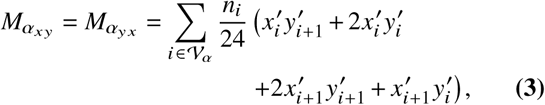

where the primed coordinates represent the translation transformation, 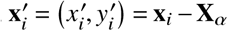, and 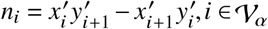. Given a rest shape matrix, 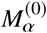, we will define cell strain as the normalized difference between both matrices,

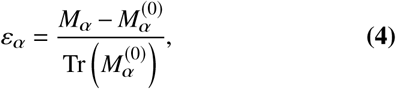

and stress with *σ*_*α*_ = *E*_*α*_*ε*_*α*_. Having described the tissue mechanically (Fig. 3C), we define the energy for a single cell as the sum of work done by turgor pressure and elastic deformation energy, resulting in the tissue mechanical energy,

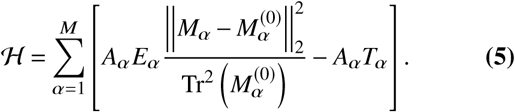

Using this model, we obtain the shape of the tissue by minimizing ℋ with respect to vertex positions.

After minimizing (equation 5), we quantify the stress acting on each wall through the average strain acting on each cell given by (equation 4). Assuming that cell wall rest length is the same between two adjacent walls compartments then it follows that they are under the same longitudinal strain, which is, to first approximation, the average between the two cells surrounding them. Let the longitudinal average strain acting on a specific wall be 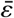. Then the stresses acting on each compartment are, by the constitutive equation, 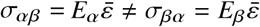. Note, that we are only considering the longitudinal components with regards to the cell wall, which means that 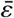 and *σ*_*α β*_ are scalar quantities. More details on the mechanical model used can be found in the supporting text.

### Tissue Mechanics - Averaged stress approximation

To assess the impact of collective mechanical behaviour within a tissue on auxin pattern self-organization we approximate the full mechanical model to a static tissue geometry where mechanical stress is averaged out across the tissue. To this end, we approximate the effects of turgor pressure of each individual cells in the static tissue by a constant average stress 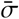 acting on it (9). The individual stress acting on a cell’s wall within the static tissue then follows directly from the stiffness of adjacent cell wall components. Again assuming that both wall compartments have the same rest length, we infer that the stress acting on a particular wall depends only on 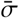 and the stiffness of the adjacent cells. Effectively, the average longitudinal strain acting on a wall surrounded by cells *α* and *β* would simply be 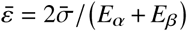. This way, instead of minimizing the full mechanical model (equation 5) given a set of turgor pressures *T*_*α*_ and rest shape matrices 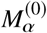 we can, in the static tissue, immediately compute stress with,

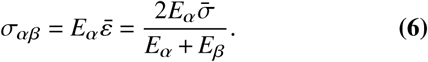

In order to quantitatively compare both methods, however, the turgor pressure, *T*_*α*_, and the rest shape matrix, 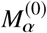, need to be chosen carefully. For an isotropic tissue (regular hexagonal lattice) in the absence of auxin patterns, all wall stresses, areas and shape matrices are the same. We can then adjust turgor pressure and rest shape matrices in the vertex model such that cell shapes and wall stresses are the same in both models, for all cells and walls.

### Auxin Transport – Compartment Model

Compartment models for auxin transport are well adapted to the context of plant development, since the prerequisite of a boundary of a plant cell is particularly well defined by courtesy of the cell wall.

Although passive diffusion occurs across cell walls, the dominant players in auxin transport are membrane-bound carriers (20, 22). Namely, efflux transporters of the PIN family are important due to their anisotropic positioning around a cell (15), which leads to a net auxin flow from one cell to the next. Let *a*_*α*_ denote an non-dimensional and normalized average auxin concentration inside cell *α*. Following the model by (9), which is similar to previous mathematical models (23, 24, 27), auxin evolves according to auxin metabolism in the cell, passive diffusion between cells and active transport across cell walls via PIN,

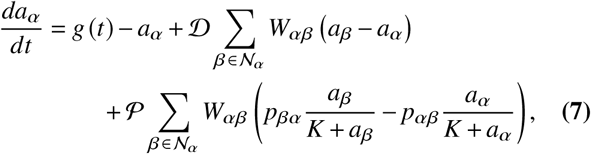

where *g* (*t*) = max {1 + *ζ* (*t*), 0} and *ζ* (*t*) a Gaussian random variable, *W*_*α β*_ = *l*_*αβ*_/*A*_*α*_, with *K, 𝒫*, and *𝒟* as adjustable parameters. 𝒟 is the magnitude of diffusion, whereas *𝒫* is the strength of PIN-mediated transport of auxin, and *K* is the Michaelis-Menten constant for the efflux of auxin. The use of a stochastic auxin expression, whose time average is ⟨*g* (*t*)⟩ ≈ 1 enables us to get closer to a biological system and will be what causes patterns to emerge. More information on how this expression is achieved can be found in the supporting text. Although this description ignores the auxin present within the extracellular domain and inside the cell wall, it has been shown that under physiological assumptions, this is a valid approximation (27). The active transport term depends on the amount of bound PIN in each cell wall,

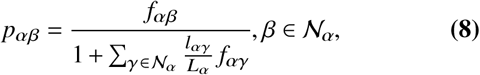

where *f*_*αβ*_, *β* ∈ *𝒩*_*α*_ expresses the ratio between binding and unbinding rates of a particular wall (Fig. 3D). Note, that *p*_*αβ*_ is different from wall to wall and from cell to cell. This means that in general, *p*_*αβ*_ ≠ *p*_*βα*_, or equivalently, *p*_*ij*_ ≠ *p*_*ji*_. This is consistent with the fact that there are two compartments to a cell wall shared by two adjacent cells. Expression (equation 8) is based on the assumption that cell walls around a particular cell compete for the same pool of PIN molecules and that the amount of PIN scales with cell perimeter. This competition has been shown to be important in the polarization of PIN (27). Alternatively, one could also scale the amount of PIN with cell size or not scale it at all. In the former case, smaller cells would be slightly preferred for auxin accumulation, whereas in the latter, larger cells would be preferred instead. Since we want to study the impact of stress patterns on the tissue, we want to decouple it from this effect as much as possible, choosing instead to scale the amount of PIN with perimeter.

The trivial fixed point of these dynamical equations is given by *a*_*α*_ = 1, ∀ *α*, which also results in equal PIN density across all walls, provided turgor pressure *T*_*α*_ and stiffness *E*_*α*_ are the same across the tissue.

The feedback between tissue mechanics and auxin pattern unfolds as auxin transport affects tissue mechanics due to auxin, *a*_*α*_, controlling cell wall stiffness, *E*_*α*_, and in reverse tissue stress, *σ*_*α*_, affects auxin transport by regulating PIN binding rates, *f*_*αβ*_, as hypothesized by (9, 10).

### Auxin-mediated cell wall softening

Auxin affects the mechanical properties of a cell wall via methyl-esterification of pectin (6, 7), resulting in a decrease of the stiffness of the cell wall. We assume that all cell wall compartments surrounding cell *α* share the same stiffness, *E*_*α*_. To capture this effect, we model stiffness with a Hill function (9) (see supporting text for more details),

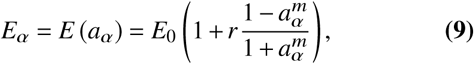

where *r*∈[0, 1[is the cell wall loosening effect, *m* is the Hill exponent of this interaction, and *E*_0_ is the stiffness of the cell walls when its auxin concentration is *a*_*α*_ = 1. At low values of auxin, *E*_*α*_ approaches the value (1 + *r*) *E*_0_, whereas at high auxin concentration, *E*_*α*_ approaches (1 − *r*) *E*_0_. Given a distribution of auxin, we can compute the wall stiffness in (equation 5) from (equation 9), or the stress acting on a specific compartment in (equation 6) for the approximated model. Here, the parameter *r* is especially important since it is directly involved in the mechanical response of the tissue when far away from the trivial steady state of auxin concentration.

### Mechanical regulation of PIN binding

According to the hypothesis presented by (9, 10), mechanical stress upregulates PIN binding. Following the model presented by (9), we model the binding-unbinding ratio, *f*_*αβ*_, as being a power law on positive stress,

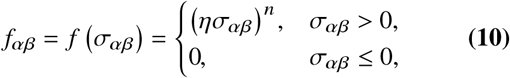

where the stresses, *σ*_*αβ*_, follow from tissue mechanics after minimization of the full mechanical model (equation 5), or, in the averaged stress approximation, it is the stress load on that particular compartment given by (equation 6). Furthermore, *n* is the exponent of this power law, and *η* captures the coupling between stress and PIN. Effectively, this mechanical coupling to PIN parameter corresponds to the sensing and subsequent response to stress, loosely translating into how much resources the cell needs to spend for processing stress cues.

### Integrating auxin transport and tissue mechanics

At each time step, Δ*t* = 0.001, starting from an auxin distribution, we compute the stiffness of each cell according to (equation 9). Then, with the input of all turgor pressures, we minimize (equation 5) to obtain tissue geometry and stresses acting on each wall. Auxin concentration in each cell will evolve according to (equation 7), where the active transport term will be regulated by stress according to (equation 10) via (equation 8). A new auxin distribution will result at the end of this iteration and we will be ready to take another time step (Fig. 4). We repeat this process until *t* = *t*_max_, where we chose *t*_max_ = 1000. Given that the patterns emerge at around *t* = 100, this choice ensures a very high likelihood that a steady state has been reached.

**Fig. 4.**
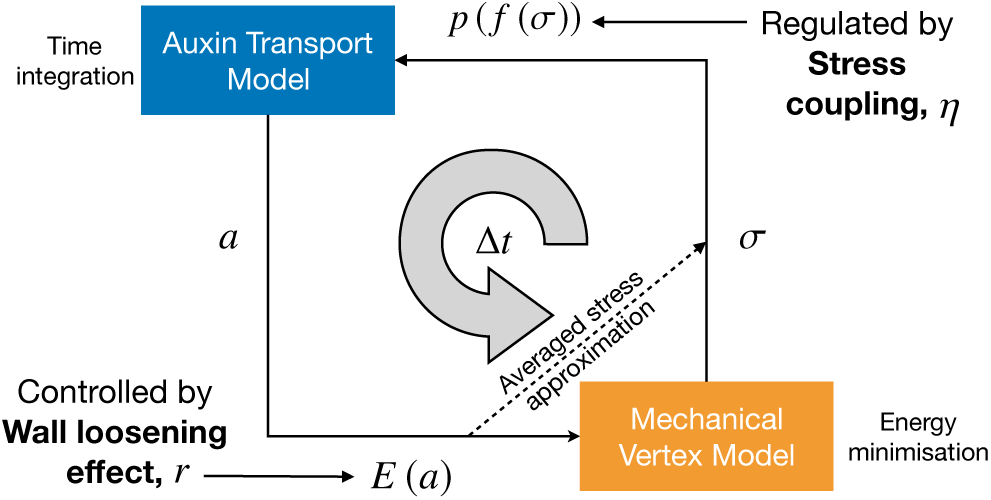
Schematic representation of the time evolution of the model. From mechanical relaxation of the mechanical model we calculate PIN densities on each wall via stress. Then we integrate auxin dynamics for a time step and update the stiffness of each cell. This process knocks the system out of the previous mechanical energy minimum and it has to be relaxed again. Alternatively, we can shortcut energy minimization using the averaged stress approximation for a static tissue. This procedure is repeated until *t* = *t*_max_. The parameters *r*, wall loosening effect, and *η*, stress coupling, interface both models and are, therefore, of critical importance to the mechanism studied.

### Implementation

We implemented this model with C++ programming language, where we have used the quad-edge data structure for geometry and topology of the tissue (52), implemented in the library QuadEdge (53). In order to minimize the mechanical energy of the tissue we have used a limited-memory Broyden-Fletcher-Goldfarb-Shanno algorithm (L-BFGS) (54, 55), implemented in the library NLopt (56). For solving the set of ODEs presented in the compartment model we used the explicit embedded Runge-Kutta-Fehlberg method (often referred to as RKF45) implemented in the GNU Scientific Library (GSL) (57). We wrapped the resulting classes into a python module with SWIG. The parameters used can be found in Table 1 of the supporting text.

### Observables

In order to quantify the existence of auxin patterns, we compute the difference between an emerging auxin concentration pattern and the trivial steady state of uniform auxin concentration pattern defined as *a*_*α*_ = 1, ∀_*α*_. To account for a large range of orders of magnitude of auxin concentration we consider as an order parameter,

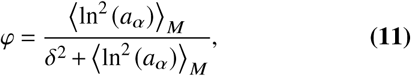

where ⟨·⟩_*M*_ denotes an average over all cells within the tissue. This way, *φ* ≈ 0 means that there are no discernible patterns, whereas *φ* ≈ 1 implies prominent auxin patterning. The term *δ*^2^ defines the sensitivity of this measure, such that an average deviation of *δ* yields *φ* ≈ 1/2 (for small *δ*). We will choose *δ* = 0.1, i.e., a 10% deviation from the trivial steady state. Since cell behaviour is tied to very high or very low auxin concentrations, we will also track the global maximum of auxin to quantify how easy it is to reach biologically relevant auxin concentrations.

Furthermore, to characterize cells with regards to PIN localization we introduce the magnitude of the average PIN efflux direction,

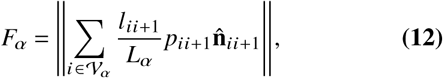

where 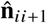 is the unit vector normal to the wall pointing outwards from *α*.

Aside from a global measure of auxin patterning, it is also important to locally relate auxin to tissue mechanics. Namely, for auxin we are interested in auxin concentration, *a*_*α*_, and auxin local gradient, obtained by interpolation,

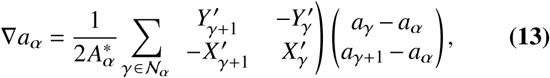

where 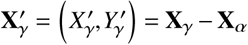 and

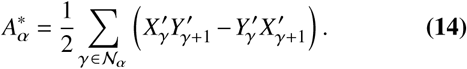

In fact, the quantity |∇*a*_*α*_| can be used as an indicator of whether there is an interface between auxin spots and the rest of the tissue.

With regards to tissue mechanics, the local quantities we quantify are the isotropic component of stress,

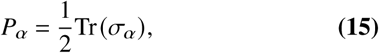

and the stress deviator tensor projected along the direction of the auxin gradient,

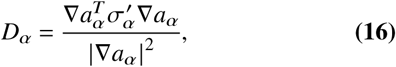

where 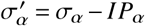, and *I* is the identity matrix. Therefore, *P*_*α*_ is a measure if a cell is being compressed (*P*_*α*_ < 0), or pulled apart (*P*_*α*_ > 0), and *D*_*α*_ translates into if a cell is more compressed along the auxin gradient than perpendicular to it (*D*_*α*_ < 0), or vice-versa (*D*_*α*_ > 0).

## Results

### The hybrid models captures stress patterns after ablation

First we verify that our hybrid model captures the expected mechanical behaviour and auxin patterning when a cell is ablated. To model ablation we set the stiffness of the ablated cell walls to *E*_0_ = 0, block all auxin transport to and from it, block PIN transporters of adjacent cells from binding to the shared wall with the ablated cell, and, finally, we lower the turgor pressure to only 10% of the original value. This remnant of pressure represents the turgor from underlying layers. This is necessary since the model only simulates the epidermal layer.

We observe that the region neighbouring the ablation site gets depleted of auxin due to PIN binding preferentially to the walls circumferentially aligned around the ablated cell (Fig. 5A). This direction lies parallel to the principal direction of stress (Fig. 5B). This stress pattern is in agreement with calculations performed by (39) in this setting and PIN aligns according to the ablation experiments in (9).

**Fig. 5.**
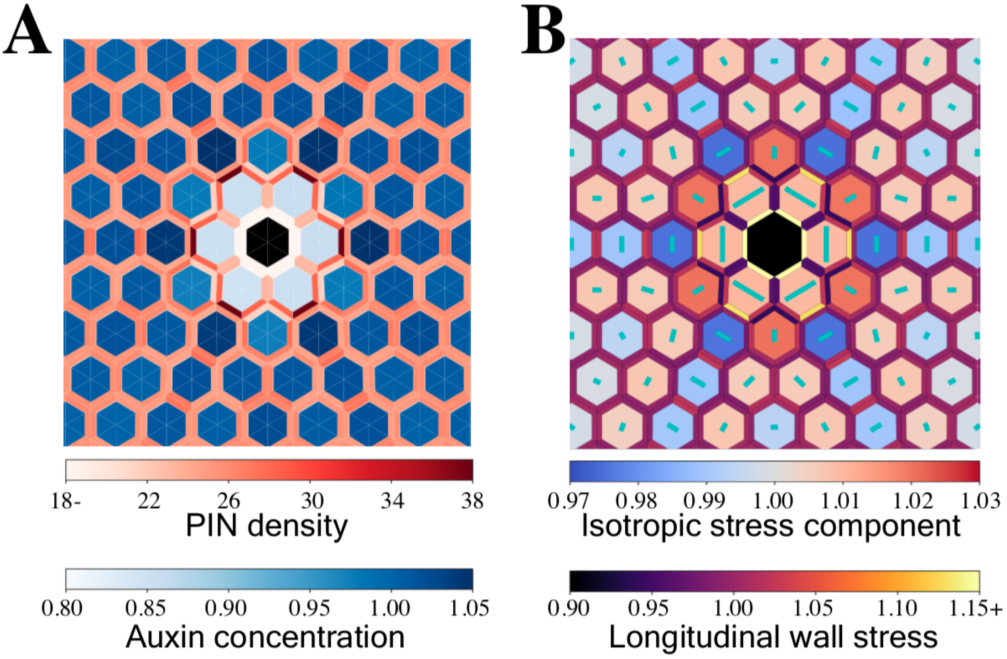
Proposed hybrid model captures expected stress patterns after ablation, as well as conditions for pattern emergence. (A) Auxin distribution, PIN density, and (B) Stress patterns as result of ablation. The green lines are directed along the largest principal direction of stress and length given by the difference of principal stresses. Ablation perturbs auxin patterning by redirecting PIN. This PIN reorientation coincides with the circumferential stress patterns around the ablation site, as seen in experiments and simulations (9, 39).

Thus our mechanical model faithfully capture the typical tissue behaviour upon ablation with regards to stress, auxin and PIN transporter patterns.

### Conditions for auxin patterns emergence

The averaged stress approximation allows to analytically compute the conditions for spontaneous auxin pattern emergence in a general regular lattice (Fig. 6). Effectively, following the analysis of (9, 27) for a regular grid, the condition for pattern formation is,

**Fig. 6.**
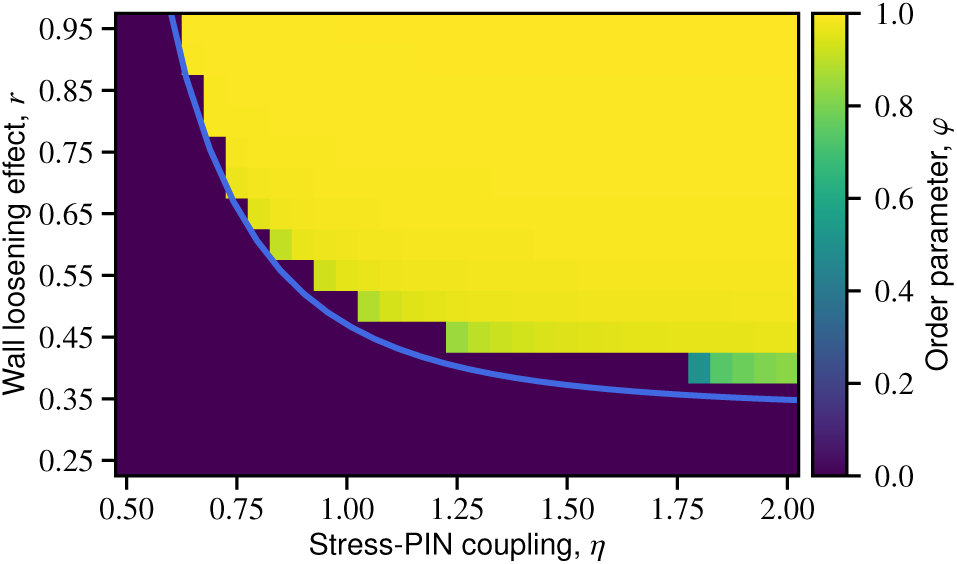
Simulation results of the order parameter *φ*, an indicator for the existence of auxin patterns, as a function of *r* ∈ [0.25, 0.95] and *η* ∈ 0.50, 2.00 for a model with tissue-wide stress patterning. The simulated tissue is composed of 2977 initially hexagonal cells. The blue line represents the analytically predicted instability for the averaged stress approximation (equation 17).

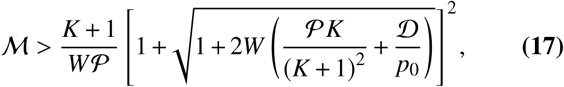

where ℳ = *nmr*, 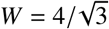 is a geometrical factor specific to the used grid, and 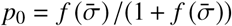 (see Supporting Text for more details).

To quantify the existence of auxin patterns in the model with tissue-wide stress patterning, we computed the normalized auxin concentration differences (order parameter *φ* defined in equation 11) for simulations with different values of wall loosening effect *r* and stress coupling *η* (Fig. 6). These two parameters are the key parameters controlling the strength of the feedback between auxin transport and cell wall mechanics.

We observe a very good agreement between the conditions for pattern emergence (equation 17) analytically predicted in the case of the averaged stress approximation and the transition of *φ* in the case of tissue-wide stress patterning (Fig. 6). This means that at the onset of patterns emergence the auxin concentrations are similar enough to make the assumption that the effect of turgor pressure is simply an isotropic stress across the entire tissue, validating the approximation near the transition. This observation is in agreement with the auxin pattern emergence mechanism hypothesis by (9) (Fig. 1).

This good agreement between the two models, however, does not necessarily apply after patterns emerge.

This poses the question of how auxin flows behave in the regime where stress patterns are also present.

### Global mechanical response reinforces PIN polarity

To understand the role of tissue-wide stress patterning on the emergence of PIN-driven auxin patterns, we quantify how PIN rearranges in the model with tissue-wide stress patterning *versus* the averaged stress approximation.

We compute the average PIN efflux direction, i.e., average PIN polarity for each combination of the critical parameters *r* (auxin induced cell wall loosening) and *η* (coupling of PIN to stress) under the averaged (Fig. 7A) and tissue-wide (Fig. 7B) stress coupling regimes.

**Fig. 7.**
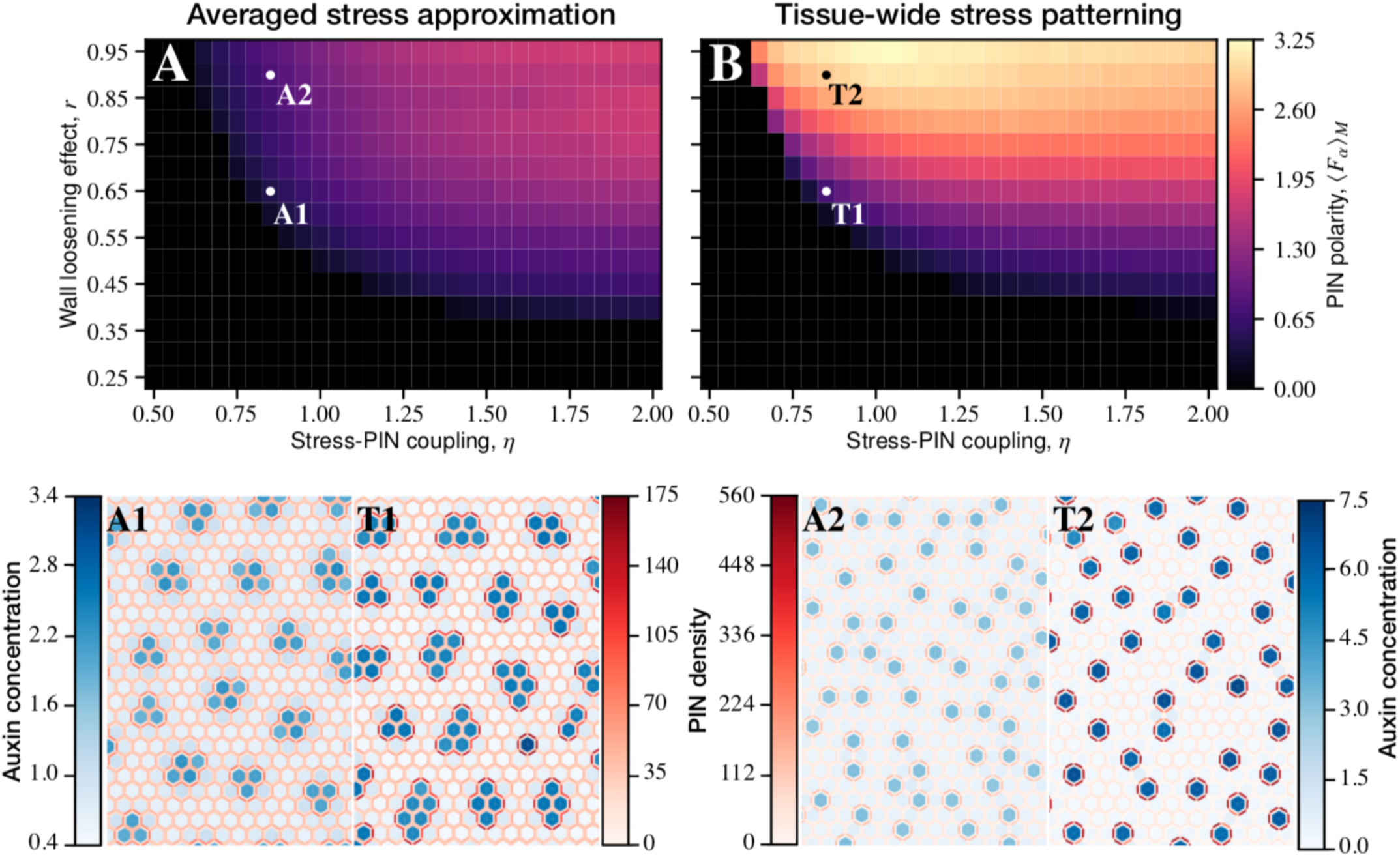
Quantification of PIN polarity. (A,B) Average magnitude of PIN polarity, ⟨*F*_*α*_ ⟩ _*M*_, as a function of stress-PIN coupling, *η*, and wall loosening effect, *r*. PIN polarity magnitude increases when considering the mechanics of the whole tissue, with a particularly strong dependence on the wall loosening affect *r* of auxin. (P1,T1,P2,T2) Comparison between example results of auxin concentration and PIN density of simulations using the averaged stress approximation (P1,P2) and the tissue-wide stress patterning (T1,T2), for the same value of *η* = 0.85, and *r* = 0.65 (P1,T1) or *r* = 0.90 (P2,T2). In both instances we observe that PIN polarity and auxin concentration are higher upon tissue-wide stress patterning (T1,T2).

We observe an overall increase in PIN polarity in the tissue-wide stress coupling regime compared with the averaged approximation. PIN polarity also becomes heavily dependent on *r*. For very low values of *r* tissue stress patterns are slightly detrimental to auxin patterning. These data show that saturation of PIN polarity happens earlier with respect to *η* for intermediate values of *r*. For high values of *r* we observe a non-monotonic dependence of polarity on *η*, effectively translating into an optimal value of *η*.

Visual inspection of the simulations results reveals higher PIN density in proximity of auxin spots and an increase in magnitude of these auxin peaks upon tissue-wide stress patterning (Fig. 7A1,T1,A2,T2). Moreover, upon tissue-wide stress patterning, cells belonging to the same auxin spot have more homogeneous auxin concentrations when comparing with the inner structure found in the auxin spots of the averaged stress approximation (Fig. 7A1,T1).

These results show that tissue-wide stress patterning reinforces PIN polarity and auxin spot concentration.

### Tissue-wide coupling induces efficient emergence of auxin spots by self-organizing stress patterns

Local auxin concentration has a profound impact on plant cell behaviour, inducing change in their mechanical properties. We thus wondered whether the reinforcement of auxin spot concentration upon tissue-wide stress patterning could in turn induce consequences on auxin patterning.

For this, we first characterize quantitatively the concentration of auxin spot measured for each simulation upon averaged stress (Fig. 8A) and tissue-wide stress (Fig. 8B) regimes. We use, as a proxy, the average of cells with auxin concentration *a*_*α*_ > 1 to identify auxin spots. We observe, that the dependence on the parameter *r* recognized for PIN polarity translates into auxin spot concentration. For medium to high values of *r*, auxin concentration is several times higher when accounting for tissue-wide behaviour than with simple cell wall stress load division.

**Fig. 8.**
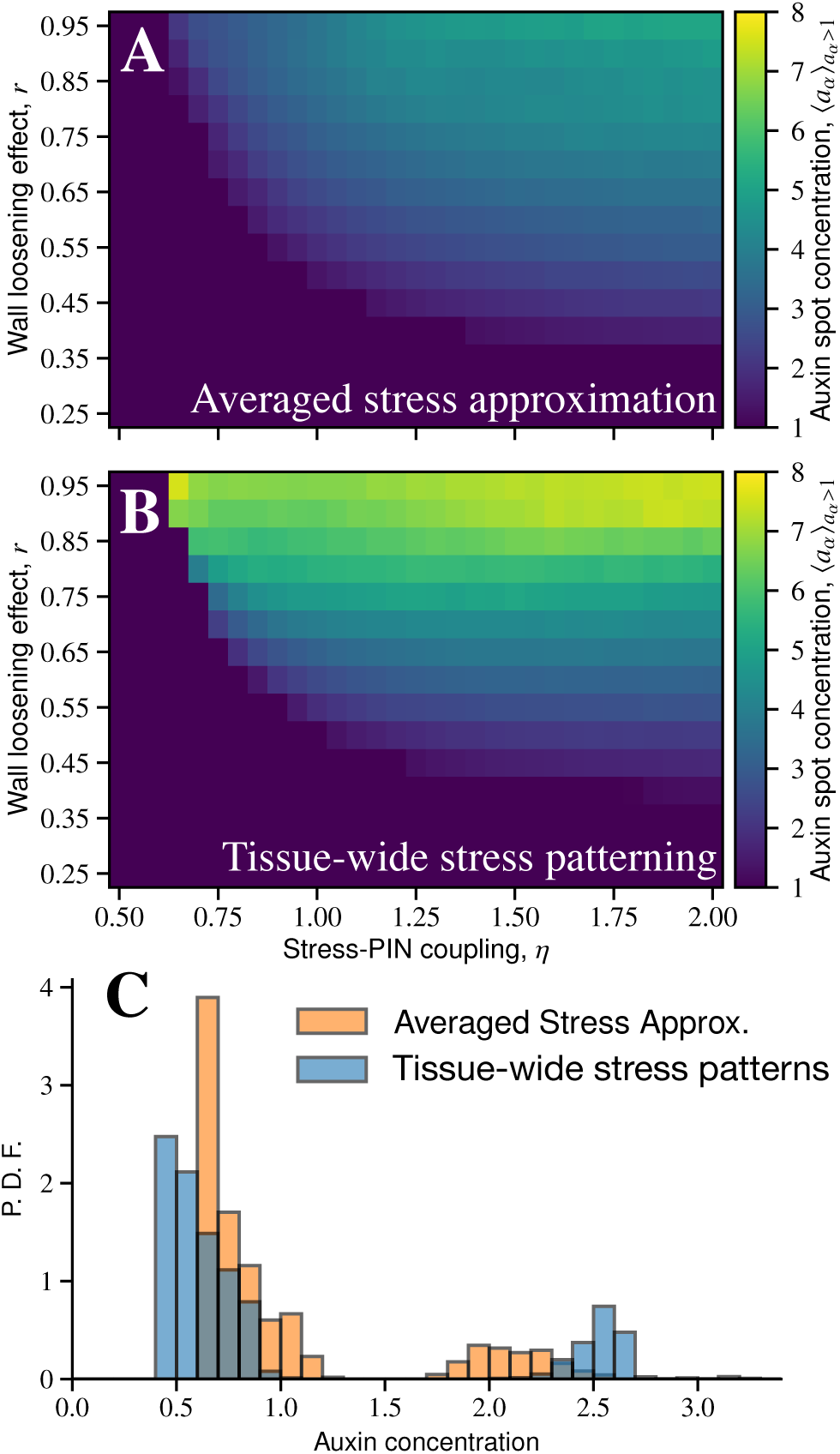
Characterization of auxin spot concentration. (A,B) Average auxin concentration for cells above basal auxin concentration (*a*_*α*_ > 1), for the averaged stress approximation (A), and upon tissue-wide stress patterning (B), as a function of stress-PIN coupling, *η*, and wall loosening effect, *r*. Spot auxin concentration increases with both *η* and *r* in (A), however, in (B), it increases predominantly with *r*. For medium to high values of *r* auxin concentration jumps to several times what would be expected, immediately after emergence. (C) For the example *r* = 0.65, *η* = 0.85, we plot the auxin distribution probability density function numerically obtained for both cases. Not only is the increase in auxin spot concentration easily discernible, but we also observe a narrower peak upon tissue-wide stress patterning, suggesting that we do indeed observe greater homogenization within the same spots, as previously shown in the examples (Fig. 7P1,T1).

Interestingly, upon tissue-wide stress patterning the concentration of auxin in peaks is poorly sensitive to *η*. Since *η* is tied to how each cell perceives wall stress, this poor sensitivity suggests that tissue-wide mechanical stress patterning feeds back into the effect of the local coupling between stress and PIN polarisation in the emergence of auxin spots. This is well illustrated when examining the distribution of auxin for a simulation with *r* = 0.65 and *η* = 0.85 (Fig. 8C. We find that the auxin spot concentration distribution is narrower upon tissue-wide stress patterning as opposed to the averaged stress approximation.

Taken together, these data suggest that biologically significant auxin patterning is achieved more efficiently upon tissue-wide stress patterning. We then asked how tissue-wide stress patterning could locally lead to more homogeneous auxin concentration. Given the link between PIN and mechanical stress, we investigated the local stress patterns around emerging auxin spots. For this, we take a close look at the results of the vertex model. For convenience, we choose an example that (a) has simple auxin patterns that allow for a straightforward interpretation, and (b) has high enough value of *r* such that tissue deformation can be easily observed. Under these conditions, we choose the parameters *r* = 0.90 and *η* = 0.85 already presented in Fig. 7T2.

To characterize the local distribution of auxin we consider both its concentration (Fig. 9A) and gradient (Fig. 9B). For the stress we consider its isotropic component (equation 15), *P*_*α*_, and the projection of deviator stress along the direction of auxin gradient (equation 16), *D*_*α*_.

**Fig. 9.**
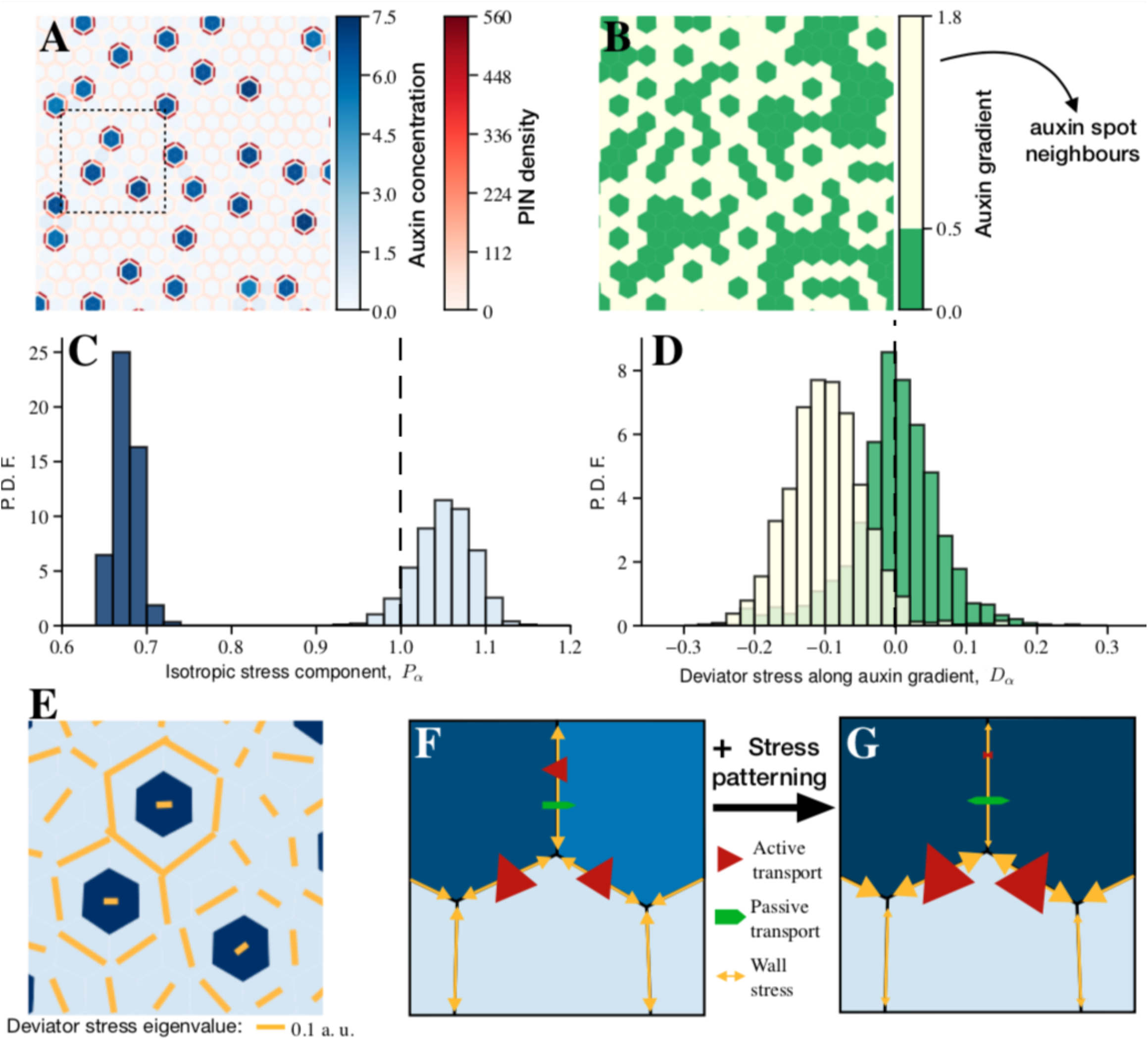
Stress self-organization according to auxin patterns. (A) Simulation result shown in Fig. 7T2 quantified here with regards to high or low auxin concentration within cells. (B) Auxin gradient categorization of cells in (A). High auxin gradient cells (> 0.5, light yellow) are neighbours of auxin spikes. (C) Distribution of the isotropic component of stress (equation 15) for high auxin concentration (dark blue), *a*_*α*_ > 3, and low auxin concentration (light blue), *a*_*α*_ ≤ 3, categories. Even though cells with larger auxin concentration expand, they are still smaller than they would be if not constrained within a tissue. Conversely, low auxin cells are affected in the opposite way. (D) Distribution of deviator stress projection along auxin direction as defined in (equation 16) for high auxin gradient cells (light yellow) and low auxin gradient cells (green). For cells adjacent to auxin spots (high gradient), the principal direction of deviator stress is perpendicular to the auxin gradient (negative value). This trend is not observed for cells with low auxin gradient category (green). The tail in the negative values for these cells results from outliers which neighbour two auxin spots. The vertical dashed lines in (C) and (D) correspond to the averaged stress approximation values. (E) Close-up taken from (A) (dashed rectangle) of deviator stress patterns, where the cells are categorized as high auxin (dark blue) and low auxin (light blue). The deviator stresses are circumferentially aligned around auxin spots due to cell stiffness differences induced by auxin cell wall softening. (F,G) Schematic representation of how stress patterning affects wall stress and, consequently, auxin transport. Circumferential stress around auxin patterns followed by PIN competition promotes PIN binding to walls perpendicular to auxin gradients at the expense of the remaining walls.

First, we classify cells into two categories, high auxin concentration (*a*_*α*_ > 3) and low auxin concentration (*a*_*α*_ ≤ 3), and compare the distribution of isotropic component of stress, *P*_*α*_ (Fig 9C). We observe that for high auxin concentration cells stress *P*_*α*_ is lower than it would be if not constrained by the remaining tissue (*P*_*α*_ = 1). Similarly, low auxin concentration cells are under larger stresses, since they are forced to accommodate the expansion of high auxin concentration cells.

Next, we looked at the direction and magnitude of stress and how it relates to the auxin gradient. We compute the projection of deviator stress onto the direction of auxin gradient (*D*_*α*_, Fig. 9D) for cells with high and low auxin gradient. The distinction into the high and low gradient categories is made because the quantity *D*_*α*_ suffers from ambiguity when auxin gradients are low. We observe a shift in the distribution of deviator stress projection *D*_*α*_ towards negative values for cells with high auxin gradients (*i*.*e*. around auxin spots) (Fig. 9D). This indicates that for cells at the interface of an auxin spot, the direction of the positive contribution to the deviator stress tensor is perpendicular to the auxin gradient.

Taken together, these data suggests that tissue deformation resulting from auxin spots patterns leads to a polarisation of stress distribution in cells around the spot: cell walls at the interface of a spot are under a larger amount of stress whereas the radial ones have decreased stress. This leads to reinforced polar auxin transport toward the spot and hence higher auxin concentration. The lower isotropic stress component inside the auxin spot furthermore supports the idea that active transport is hampered between cells with high auxin concentration, leading to the observed homogenisation of auxin spots (Fig. 9F,G). Altogether these data show that tissue-wide stress patterning efficiently promotes emergence of auxin spots by locally self-organising stress patterns and PIN polarisation.

## Discussion

Here, we used a hybrid model composed of a vertex model for plant tissue mechanics, and a compartment model for auxin transport to uncover the role of tissue-wide mechanical coupling on auxin redistribution. We first verify that our model successfully captures the behaviour of plant tissue upon ablation experiments and the conditions for emergence of auxin patterns. We then compared the behaviour of our model featuring tissue-wide mechanical coupling to an approximation which averages out stress across the tissue. We observe the emergence of focused auxin spots with high auxin concentration when tissue-wide mechanical coupling is implemented, not when stress is averaged. We show that tissue-wide mechanical coupling favours the emergence of anisotropic circumferential stress field around the auxin spots, which reinforces PIN polarity and homogenizes auxin concentrations within the auxin spots. Emerging tissue-wide stress patterns therefore increase the efficiency of auxin patterns emergence by mechanical polarisation of PIN in response to local circumferential stress field around the spots.

The auxin-induced cell wall loosening effect (*r* parameter in this work) is an important determinant of the feedback of auxin on the tissue mechanics. The range of values of *r* for which substantial pattern focusing occurs is around *r* ∼ 0.60 and above in our model. This translates into a variation of stiffness from a minimum value *E*_min_ up to *E*_max_ = 4*E*_min_ (see supplementary material). Although high, this range is within biological expectation and supported by AFM measurements on auxin treated tissues (7) and comparable to previous simulations of this mechanism (9) where *E*_max_/*E*_min_ = 5 which translates into *r* = 2/3, a value in the self-reinforcing region. Furthermore, since plants create auxin patterns in specific regions, it stands to reason that plant tissue can control *η* within a range that includes the transition. Therefore, our choice of range for *η* is a reasonable one.

Comparison of the tissue-wide stress patterning case to the averaged stress approximation reveals that auxin spot concentration has a very steep transition in the former case (Fig. 8A,B). This results in a several-fold increase in auxin concentration at values of stress-PIN coupling (*η*, the stress detection parameter) barely above threshold for pattern formation. This leads to the appearance of discernible auxin spots without requiring a sensitive stress detection. Tissue-wide stress patterning, as it results from tissue relaxation, improves the efficiency of auxin patterning at no extra cost to the plant. The coupling serves as an amplifier of mechanical cues which sharpen locally auxin patterns. This mechanism is inherently efficient as low values of mechanical cue detection the patterns can overcome noise and external factors contributing to robustness.

How is this local reinforcement of auxin pattern induced by tissue-wide mechanical coupling achieved? Our results show that circumferential stress patterns promote polarisation of PIN toward the spots feed forwarding auxin into the spots. The emerging circumferential stress patterns caused by emerging auxin spots and those in the ablation simulation (Fig.5B) are very similar. This is of no coincidence, since both entail a local decrease in stiffness. Remarkably, however, is that this circumferential stress pattern also coincides with the shape-induced stress patterns, as indicated by microtubule orientation, around the tip of the primordium as it emerges from the meristem (39). Therefore, tissue-wide stress patterning set the stage for primordium outgrowth by focusing efficiently auxin, forming local circumferential stress that in turn may re-orient microtubules and prefigure the shape of the primordium. This process could, in turn, be capable of reinforcing auxin transport to the tip of the newly forming organ. Yet, quantifying this requires further modeling. Another consequence of tissue-wide mechanical coupling revealed here, is the homogenization of auxin concentration inside a spot. An implication could be that an homogeneous auxin concentration facilitates the primordium bulging out from the meristem. It would be important to link our observations to simulations of the primordium outgrowth as presented in the work of (8).

## Conclusion

Even though the mechanisms by which PIN preferentially associate with stressed cell walls is unclear, here we show that there are substantial advantages by intertwining tissue-wide mechanics and auxin patterning. Even if auxin patterning is possible by chemical processes and local mechanical coupling, tissue-wide mechanics may provide a way for patterning to still occur at a lower energy cost for the tissue. Moreover, this process can also provide robustness to the patterning, factoring in tissue-wide stress pattern, a sort of proprioceptive mechanism.

## ACKNOWLEDGEMENTS

This work was supported by the Max Planck Society and the Deutsche Forschungsgemeinschaft via DFG-FOR2581.

## Competing interests

The authors declare no conflicts of interest.

