## Supplemental Text for "Tissue-wide integration of mechanical cues promotes efficient auxin patterning"

### Derivation of Auxin Transport Model

Geometrically, we consider the epithelial tissue to be composed of cells shaped as right prisms with the same height $h$. Effectively, under this assumption we can reduce the dimensionality of the system by considering just the shape of the base of the cells. We will also assume that the thickness of the walls is thin enough such that the volume of the cell is approximately $hA_\alpha$, where $A_\alpha$ is the area of the base of the cell. Let $\rho$ denote auxin concentration. Assuming auxin gets expressed at a rate $\gamma$ per unit time and unit volume, and each auxin molecule decays at a rate $\delta$ per unit time, we can write the continuity equation inside any cell as

$$\dot{\rho} + \nabla \cdot \mathbf{J} = \gamma - \delta\rho. \tag{1}$$

where $\mathbf{J}$ is local current density of auxin.
Focusing on cell $\alpha$, we can integrate inside the cell to obtain,

$$\dot{N}_\alpha + \oiint_{S_\alpha} \mathbf{J} \cdot \hat{\mathbf{n}} dS = hA_\alpha\gamma - \delta N_\alpha, \tag{2}$$

where we assume the volume $V_\alpha$ does not change over time, $S_\alpha$ is the surface enclosing volume $V_\alpha$, and $N_\alpha$ represents the number of auxin molecules inside cell $\alpha$. Another assumption we make is that the flux of $\mathbf{J}$ across the bases of the cell is zero, resulting in just flux across neighbours whithin the same layer.
To simplify notation we define $\mathcal{N}_\alpha$ as the set of all regions on the other side of each wall surrounding cell $\alpha$. This set will contain the neighbouring cells. Note that with this definintion there can be multiple occurrences of the same cell within the set, yet referred to as different regions. We can now uniquely refer to wall compartments around a cell with $\alpha\beta$ or $\beta\alpha$, $\beta \in \mathcal{N}_\alpha$.
If we let the wall between cells $\alpha$ and $\beta \in \mathcal{N}_\alpha$ be of length $l_{\alpha\beta}$ with normal vector $\hat{\mathbf{n}}_{\alpha\beta}$ pointing from $\alpha$ to $\beta$, then we can replace the surface integral by a sum of line integrals,

$$\dot{N}_\alpha + h \sum_{\beta \in \mathcal{N}_\alpha} \int_{l_{\alpha\beta}} \mathbf{J}(\mathbf{r}) \cdot \hat{\mathbf{n}}_{\alpha\beta} dl = hA_\alpha\gamma - \delta N_\alpha. \tag{3}$$

Assuming auxin diffuses quickly inside each cell, we can assume auxin concentration is the same everywhere inside the cell, $\rho_\alpha = N_\alpha/(hA_\alpha)$. This approximation is justified, since auxin is a very small molecule. Now, with a constant auxin concentration whithin a cell, we can write the passive transport with Fick's law, i.e., $\mathbf{J} = \mathbf{J}_{\text{pass}} + \mathbf{J}_{\text{act}}$, with $\mathbf{J}_{\text{pass}} = -D\nabla\rho$ and interpolating auxin concnetration inside the cell wall. Therefore, the current density of auxin through wall $l_{\alpha\beta}$ from cell $\alpha$ to region $\beta$ is

$$\mathbf{J}_{\text{pass}_{\alpha\to\beta}} \cdot \hat{\mathbf{n}}_{\alpha\beta} = -D\frac{\rho_\beta - \rho_\alpha}{T}, \tag{4}$$

where $T$ is the thickness of the wall. We can also continue to simplify assuming that the active auxin flow is constant along the wall. Again, assuming that the area $A_\alpha$ does not change,

$$\dot{\rho}_\alpha = \gamma - \delta\rho_\alpha + \frac{D}{T}\sum_{\beta \in \mathcal{N}_\alpha} \frac{l_{\alpha\beta}}{A_\alpha}\left(\rho_\beta - \rho_\alpha\right) + \sum_{\beta \in \mathcal{N}_\alpha} \frac{l_{\alpha\beta}}{A_\alpha}\left[J_{\text{act}_{\beta\to\alpha}} - J_{\text{act}_{\alpha\to\beta}}\right], \tag{5}$$

where $J_{\text{act}_{\alpha\to\beta}}$ is the amount of auxin transported per unit area, per unit time, from $\alpha$ to $\beta$.
Let $P_{\alpha\beta}$ be the density of auxin efflux carriers on the compartment of the wall facing cell $\alpha$, pumping auxin into $\beta$. We model active auxin transport with

$$J_{\text{act}_{\alpha\to\beta}} = \mathcal{A} P_{\alpha\beta}\frac{\rho_\alpha}{K_M + \rho_\alpha}, \tag{6}$$

with $\mathcal{A}$ being the maximum activity of the carrier and $K_M$ the Michaelis-Menten constant of the reaction. This results in

$$\dot{\rho}_\alpha = \gamma - \delta\rho_\alpha + \frac{D}{T}\sum_{\beta \in \mathcal{N}_\alpha} \frac{l_{\alpha\beta}}{A_\alpha}\left(\rho_\beta - \rho_\alpha\right) + \mathcal{A}\sum_{\beta \in \mathcal{N}_\alpha} \frac{l_{\alpha\beta}}{A_\alpha}\left[P_{\beta\alpha}\frac{\rho_\beta}{K_M + \rho_\beta} - P_{\alpha\beta}\frac{\rho_\alpha}{K_M + \rho_\alpha}\right]. \tag{7}$$

We now need to model PIN molecule binding and unbinding. Let the number of internalized PIN molecules be $M_\alpha$. Let the number of PIN molecules bound to wall $l_{\alpha\beta}$ of cell $\alpha$ pumping auxin into region $\beta$ by $M_{\alpha\beta}$. Also, we define the binding and unbinding constants, $k_{\text{b}}$ and $k_{\text{ub}}$. The rate equations for bound PIN molecules are

$$\dot{M}_{\alpha\beta} = k_{\text{b}}\frac{l_{\alpha\beta}}{L_\alpha}M_\alpha - k_{\text{ub}}M_{\alpha\beta}, \tag{8}$$

where $L_\alpha = \sum_{\beta\in\mathcal{N}_\alpha} l_{\alpha\beta}$ is the perimeter of the cell. If $k_\mathrm{b}$ takes into account how many PIN molecules are close enough to the walls of the cell, the factor $\frac{l_{\alpha\beta}}{L_\alpha}$ is necessary since it is the probability of being close to that particular wall. In terms of PIN density $P_{\alpha\beta}$ and concentration $P_\alpha$ instead of PIN number,

$$\dot{P}_{\alpha\beta} = k_\mathrm{b}\frac{A_\alpha}{L_\alpha}P_\alpha - k_\mathrm{ub}P_{\alpha\beta}, \tag{9}$$

We will consider the dynamics of PIN to be much faster than that of auxin, such that we can decouple the time scales of both processes. As such, we assume that between auxin time steps there is enough time for PIN to reach its steady state. The steady state of this equation is

$$P^*_{\alpha\beta} = \frac{k_\mathrm{b}}{k_\mathrm{ub}}\frac{A_\alpha}{L_\alpha}P^*_\alpha = f_{\alpha\beta}\frac{A_\alpha}{L_\alpha}P^*_\alpha, \tag{10}$$

where the binding and unbinding rates are a function of the features of the wall, $f_{\alpha\beta} = k_\mathrm{b}/k_\mathrm{ub}$.
Assuming all cells are competing for the same amount of PIN molecules, $M_0$, we have

$$A_\alpha P^*_\alpha + \sum_{\beta\in\mathcal{N}_\alpha} l_{\alpha\beta}P^*_{\alpha\beta} = M_0, \tag{11}$$

which, using $P^*_{\alpha\beta}$ from the previous expression, can be be solved for $P^*_\alpha$, yielding

$$P^*_\alpha = \frac{M_0}{A_\alpha}\frac{1}{1+\sum_{\beta\in\mathcal{N}_\alpha}\frac{l_{\alpha\beta}}{L_\alpha}f_{\alpha\beta}}, \tag{12}$$

and for $P^*_{\alpha\beta}$,

$$P^*_{\alpha\beta} = \frac{M_0}{L_\alpha}\frac{f_{\alpha\beta}}{1+\sum_{\xi\in\mathcal{N}_\alpha}\frac{l_{\alpha\xi}}{L_\alpha}f_{\alpha\xi}} = \frac{M_0}{L_\alpha}p_{\alpha\beta}. \tag{13}$$

Note that if we replace $P^*_{\alpha\beta}$ for $P_{\alpha\beta}$ in auxin concentration rate equation with $M_0$ being a constant, when solving for steady state, we obtain that the steady state concentration of auxin is dependent on the geometry of the cell, namely its perimeter.
There are three choices we can make, in regards to how plant cells scale PIN. We can simply not scale resulting in a constant $M_0$, we can scale according to cell area, i.e., $M_0 = A_\alpha C_0$, or scale according to perimeter, $M_0 = L_\alpha D_0$. To minimise the impact of this effect on auxin patterning, we opt for a scaling according to perimeter, resulting in the density

$$P_{\alpha\beta} = D_0 p_{\alpha\beta}. \tag{14}$$

By replacing in the auxin concentration rate equation we finally get

$$\dot{\rho}_\alpha = \gamma - \delta\rho_\alpha + \frac{D}{T}\sum_{\beta\in\mathcal{N}_\alpha}\frac{l_{\alpha\beta}}{A_\alpha}\left(\rho_\beta - \rho_\alpha\right) + \mathcal{A}D_0\sum_{\beta\in\mathcal{N}_\alpha}\frac{l_{\alpha\beta}}{A_\alpha}\left[p_{\beta\alpha}\frac{\rho_\beta}{K_M+\rho_\beta} - p_{\alpha\beta}\frac{\rho_\alpha}{K_M+\rho_\alpha}\right], \tag{15}$$

where,

$$p_{\alpha\beta} = \frac{f_{\alpha\beta}}{1+\sum_{\xi\in\mathcal{N}_\alpha}\frac{l_{\alpha\xi}}{L_\alpha}f_{\alpha\xi}} \tag{16}$$

We adimensionalize the equation by choosing a characteristic length $L$ and a characteristic time $1/f$. We can then replace, $\rho = \rho^*/L^3$, $dt = dt^*/f$, $l = Ll^*$, $K = K^*/L^3$ and $A = A^*L^2$,

$$\frac{d\rho^*_\alpha}{dt^*} = \frac{L^3\gamma}{f} - \frac{\delta}{f}\rho^*_\alpha + \frac{D}{LfT}\sum_{\beta\in\mathcal{N}_\alpha}\frac{l^*_{\alpha\beta}}{A^*_\alpha}\left(\rho^*_\beta - \rho^*_\alpha\right) + \frac{\mathcal{A}D_0L^2}{f}\sum_{\beta\in\mathcal{N}_\alpha}\frac{l^*_{\alpha\beta}}{A^*_\alpha}\left[p_{\beta\alpha}\frac{\rho^*_\beta}{K^*+\rho^*_\beta} - p_{\alpha\beta}\frac{\rho^*_\alpha}{K^*+\rho^*_\alpha}\right]. \tag{17}$$

Note that the auxin concentration in steady state, when all pin densities $p_{\alpha\beta}$ are the same, is $\rho^*_0 = L^3\gamma/\delta$. If we normalize the previous expression by choosing $\rho^*_0$ as our unit of auxin concentration and $\delta/f$ as our unit of time, we obtain

$$\frac{da_\alpha}{d\tau} = \frac{f}{L^3\gamma}\frac{d\rho^*_\alpha}{dt^*} = 1 - a_\alpha + \frac{D}{L\delta T}\sum_{\beta\in\mathcal{N}_\alpha}\frac{l^*_{\alpha\beta}}{A^*_\alpha}\left(a_\beta - a_\alpha\right) + \frac{\mathcal{A}D_0}{L\gamma}\sum_{\beta\in\mathcal{N}_\alpha}\frac{l^*_{\alpha\beta}}{A^*_\alpha}\left[p_{\beta\alpha}\frac{a_\beta}{K+a_\beta} - p_{\alpha\beta}\frac{a_\alpha}{K+a_\alpha}\right], \tag{18}$$

where $a_\alpha = \rho_\alpha^*/\rho_0^*$, $d\tau = dtf/\delta$ and $K = K^*/\rho_0^*$. The combination of parameters relevant for this process are $\mathcal{P} = \frac{\mathcal{A}D_0}{L\gamma}$ and $\mathcal{D} = \frac{D}{L\delta T}$. If the thickness of the wall is also adimensional, then $\mathcal{D} = \frac{D}{L^2\delta T^*}$. These two numbers are essentially a comparison between passive or active transport and auxin turnover rate. We can also have a stochastic expression rate in order to break symmetry and to make the system closer to a biological one. Let $g(\tau)$ be a normally distributed random variable with mean $\mu = 1$ and standard variation $\sigma$. We will also restrict to the values $g(\tau) \geq 0$. Finally,

$$\frac{da_\alpha}{d\tau} = g(\tau) - a_\alpha + \mathcal{D} \sum_{\beta\in\mathcal{N}_\alpha} \frac{l_{\alpha\beta}^*}{A_\alpha^*}\left(a_\beta - a_\alpha\right) + \mathcal{P} \sum_{\beta\in\mathcal{N}_\alpha} \frac{l_{\alpha\beta}^*}{A_\alpha^*}\left[p_{\beta\alpha}\frac{a_\beta}{K + a_\beta} - p_{\alpha\beta}\frac{a_\alpha}{K + a_\alpha}\right], \tag{19}$$

where

$$p_{\alpha\beta} = \frac{f_{\alpha\beta}}{1 + \sum_{\xi\in\mathcal{N}_\alpha}\frac{l_{\alpha\xi}}{L_\alpha} f_{\alpha\xi}}. \tag{20}$$

The connection of this model to tissue mechanics happens with modeling binding rates $f_{\alpha\beta}$ as a function of wall stress and modeling auxin-mediated cell wall loosening, by changing stiffness $E_\alpha$.
Suppose stress $\sigma_{\alpha\beta}$ is the longitudinal component of stress acting on the wall between cell $\alpha$ and region $\beta \in \mathcal{N}_\alpha$, specifically in the wall compartment facing cell $\alpha$. In accordance with [1], we model $f_{\alpha\beta}$ as a power-law,

$$f_{\alpha\beta} = f\left(\sigma_{\alpha\beta}\right) = \begin{cases} \left(\eta\sigma_{\alpha\beta}\right)^n, & \sigma_{\alpha\beta} > 0, \\ 0, & \sigma_{\alpha\beta} \leq 0. \end{cases} \tag{21}$$

Finally, we look at cell wall loosening. We consider that all compartments surrounding a specific cell have the same stiffness. We model auxin-mediated cell wall loosening with a hill function similarly to [1],

$$E_\alpha = E\left(a_\alpha\right) = E_{\max} - \left(E_{\max} - E_{\min}\right)\frac{a_\alpha^m}{1 + a_\alpha^m}, \tag{22}$$

where $m$ is the Hill exponent, $a_\alpha = 1$ $(\rho_\alpha = \rho_0)$ being the value of auxin where the stiffness is $\left(E_{\max} + E_{\min}\right)/2$, and $E_{\min}$ and $E_{\max}$ the minimum and maximum values stiffness in this system. It is also useful for linear stability analysis to rewrite this expression in terms of the parameters

$$E_0 = \left(E_{\max} + E_{\min}\right)/2, \qquad r = \frac{E_{\max} - E_{\min}}{E_{\max} + E_{\min}}. \tag{23}$$

Inverting this transformation we get,

$$E_{\max} = E_0\left(1 + r\right), \qquad E_{\min} = E_0\left(1 - r\right), \tag{24}$$

and the final expression for cell wall loosening becomes

$$E_\alpha = E\left(a_\alpha\right) = E_0\left(1 + r\frac{1 - a_\alpha^m}{1 + a_\alpha^m}\right). \tag{25}$$

### Mechanical model details

We assign a mechanical energy to the tissue as a function of vertex positions. We can then minimise the mechanical energy with respect to vertex positions to find the current tissue configuration. To simplify notation, let $\mathcal{V}_\alpha$ be a set of vertices around the polygon representing cell $\alpha$, ordered counter-clockwise. We will also have the set be cyclic, i.e., if the cell has $V$ vertices, then $i \in \mathcal{V}_\alpha$, $i = 1, 2, \ldots, V$ implies $\mathbf{x}_{i+1} = \mathbf{x}_1$ when $i = V$ and $\mathbf{x}_{i-1} = \mathbf{x}_V$ when $i = 1$. This way we can uniquely define a wall compartment between two vertices.
The two main terms we are going to focus on is the work of the turgor pressure $T_\alpha$, given by $-\int_{V_\alpha} T_\alpha dV_\alpha = -T_\alpha A_\alpha h$, and the elastic energy $\int_{V_\alpha} \psi_\alpha dV_\alpha$, where $\psi_\alpha$ is the elastic energy density $\varepsilon_\alpha : \sigma_\alpha$, where the : represents the element-wise contraction of matrices. We will use form matrices $M_\alpha$ to write down a proxy to strain as

$$\varepsilon_\alpha = \frac{M_\alpha - M_\alpha^{(0)}}{\operatorname{Tr}\left(M_\alpha^{(0)}\right)}, \tag{26}$$

where $M_\alpha^{(0)}$ is a target shape matrix and where these shape matrices are given by,

$$M_{\alpha_{xx}} = \sum_{i\in\mathcal{V}_\alpha} \frac{n_i}{12}\left(x_i'^2 + x_i' x_{i+1}' + x_{i+1}'^2\right), \tag{27}$$

$$M_{\alpha_{yy}} = \sum_{i\in\mathcal{V}_\alpha} \frac{n_i}{12}\left(y_i'^2 + y_i' y_{i+1}' + y_{i+1}'^2\right), \tag{28}$$

$$M_{\alpha_{xy}} = M_{\alpha_{yx}} = \sum_{i\in\mathcal{V}_\alpha} \frac{n_i}{24}\left(x_i' y_{i+1}' + 2x_i' y_i' + 2x_{i+1}' y_{i+1}' + x_{i+1}' y_i'\right), \tag{29}$$

where the primed coordinates represent the translation transformation, $\mathbf{x}_i' = \left(x_i', y_i'\right) = \mathbf{x}_i - \mathbf{X}_\alpha$, and $n_i = x_i' y_{i+1}' - x_{i+1}' y_i'$, $i \in \mathcal{V}_\alpha$. Also, we can easily compute the area with

$$A_\alpha = \frac{1}{2}\sum_{i\in\mathcal{V}_\alpha} n_i. \tag{30}$$

We make the approximation that the material is linear, isotropic, and uniform, which means that the constitutive equation is given by the ususal shape This means that stress under these assumptions has two free parameters, the Lamé constants $\mu$ and $\lambda$, and is given by

$$\sigma_\alpha = 2\mu\varepsilon_\alpha + \lambda I \operatorname{Tr}\left(\varepsilon_\alpha\right). \tag{31}$$

Another assumption that we make is that the Poisson ratio is small enough to disregard the second term of this expression. The factor $2\mu$ is the auxin-dependent stiffness, yielding

$$\sigma_\alpha \approx E\left(a_\alpha\right)\varepsilon_\alpha. \tag{32}$$

Now we can write the energy density as $E\left(a_\alpha\right)||\varepsilon_\alpha||^2$, where $||A||^2 = A\colon A$. Integrating over the volume of the cell and summing the elastic term and turgor pressure for all cells, we arrive at the Hamiltonian

$$\mathcal{H} = \sum_\alpha \mathcal{H}_\alpha = \sum_\alpha \left[A_\alpha E\left(a_\alpha\right)\frac{\left\|M_\alpha - M_\alpha^{(0)}\right\|^2}{\operatorname{Tr}^2\left(M_\alpha^{(0)}\right)} - A_\alpha T_\alpha\right], \tag{33}$$

where we factored out the cell height $h$ since the minimum of this expression becomes independent from it. What remains is how to extract a longitudinal stress acting on each cell wall compartment.

Consider a composite wall of length $l$ and cross section $S$ made from two compartments $A$ and $B$ in parallel with different stiffness, $E_A$ and $E_B$, under the same strain $\varepsilon$. The sum of the elastic energy in both compartments is $lSE_A\varepsilon^2/2 + lSE_B\varepsilon^2/2 = lS\bar{\sigma}\varepsilon$, i.e. $\varepsilon = 2\bar{\sigma}/(E_A + E_B)$. Therefore, $\sigma_A = E_A\varepsilon = 2E_A\bar{\sigma}/(E_A + E_B)$. Note that we can approximate the tugor pressure by prescribing a value of stress, $\bar{\sigma}$, and bypass mechanical relaxation of the tissue. We will call this the averaged stress approximation. Nevertheless, the pattern formation mechanism hinges on the assumption that the strain is the same on both compartments of the cell wall, meaning that we still have to estimate the strain a wall is under from average cell strain. In order to estimate that value, we interpolate strain between the two adjacent cells and project that value onto the wall to get the average longitudinal strain acting on that specific wall. Then, we can simply use the constitutive equation to get the longitudinal stress of each compartment based on their stiffness. Note that with the definition used for elastic energy density, a factor of 1/2 needs to be accounted for in $E_0$ in order to compare to the usual Young's modulus. Also note that since we are only looking for the minimum of this expression, we can pick any arbitrary units for $E_\alpha$ provided we perform the same conversion for $T_\alpha$ and obtain the same geometry $A_\alpha$ and $M_\alpha$.

### Algorithm and implementation details

The tissue geometry used was an hexagonal lattice in units such that the each side of each hexagon is of length $1L$. First we need the tissue in the vertex model and in the approximated model to have the same shape in the abscence of auxin patterns, $M_\alpha^{\text{init}}$, in order for the results between both approaches to be comparable. Also, in the averaged stress approximation we need to prescribe a $\bar{\sigma}$ which will correspond to a combination of tugor pressure and deformation. Therefore, we fit the values of $T_\alpha$ and $M_\alpha^{(0)}$ $(\forall\alpha)$ to the lattice size and $\bar{\sigma}$ we prescribe, in order for both models to be indistinguishable when no patterns emerge ($a_\alpha = 1$, $\forall\alpha$).

Table 1: Table of simulation parameters, their meaning and value or range. All parameters are in normalized units close to unity for numerical accuracy. The parameters $T_\alpha$ and $M_\alpha^{(0)}$ were fitted such that the initial shape of the tissue (without auxin patterns) corresponds to a regular hexagonal lattice of side 1, $M_\alpha^{\text{init}}$, and wall stress for all walls equals $1 = \bar{\sigma}$.

| Symbol | Meaning | Value or Range (a.u.) |
|---|---|---|
| $\text{Var}[g(\tau)]$ | normalized expresssion rate variance | 0.09 |
| $\mathcal{D}$ | diffusion magnitude number | 20 |
| $\mathcal{P}$ | active transport strength number | 100 |
| $K$ | Michaelis-Menten constant | 1 |
| $\eta$ | stress-PIN coupling constant | $[0.5, 2.0]$ |
| $n$ | stress-PIN coupling power | 3 |
| $r$ | wall loosening effect | $[.25, .95]$ |
| $m$ | wall loosening power | 2 |
| $E_0$ | normalized reference stiffness | 24 |
| $T_\alpha$ | normalized turgor pressure | $2.2088,\ \forall\alpha$ |
| $M_\alpha^{(0)}$ | rest shape matrix | $0.92308 M_\alpha^{\text{init}},\ \forall\alpha$ |
| $\bar{\sigma}$ | normalized average stress | 1 |
| $\Delta t$ | simulation time step | 0.001 |
| $t_{\max}$ | total simulation time | 1000 |

We assume very different timescales for auxin transport and mechanical cue propagation through the tissue. This means that there is enough time for the tissue to relax to a local minimum in between auxin transport time steps, $\Delta t$. To simulate this model, we start by computing wall stress in the current configuration, calculate PIN density on all cell walls with (20), take a step $\Delta t$ in the set of ODEs (19), with new auxin concentrations compute stiffness with (25), and, finally, minimise mechanical energy (33). This is done until the maximum simulation time $t_{\max}$ has been reached. The simulation parameters used can be found in table (1).
Computationally the minimization procedure is the performance bottleneck. In order to overcome that, we used a gradient-based minimisation method (L-BFGS) and computed the gradient of (33), which is straightforward, albeit tedious.

### Linear Stability Analysis

The auxin transport model in use is

$$\frac{da_\alpha}{d\tau} = g(\tau) - a_\alpha + \mathcal{D} \sum_{\beta \in \mathcal{N}_\alpha} \frac{l^*_{\alpha\beta}}{A^*_\alpha} \left(a_\beta - a_\alpha\right) + \mathcal{P} \sum_{\beta \in \mathcal{N}_\alpha} \frac{l^*_{\alpha\beta}}{A^*_\alpha} \left[ p_{\beta\alpha} \frac{a_\beta}{K + a_\beta} - p_{\alpha\beta} \frac{a_\alpha}{K + a_\alpha} \right], \tag{34}$$

where

$$p_{\alpha\beta} = \frac{f_{\alpha\beta}}{1 + \sum_{\xi \in \mathcal{N}_\alpha} \frac{l_{\alpha\xi}}{L_\alpha} f_{\alpha\xi}}, \tag{35}$$

$$f_{\alpha\beta} = f\left(\sigma_{\alpha\beta}\right) = \begin{cases} \left(\eta\sigma_{\alpha\beta}\right)^n, & \sigma_{\alpha\beta} > 0, \\ 0, & \sigma_{\alpha\beta} \le 0, \end{cases} \tag{36}$$

and,

$$E_\alpha = E\left(a_\alpha\right) = E_0 \left(1 + r \frac{1 - a_\alpha^m}{1 + a_\alpha^m}\right) \tag{37}$$

Consider the case when all cells have the same auxin concentration $a_\alpha = 1$. All stiffnesses are the same and therefore we can regard the effect of turgor pressure as a constant stress of magnitude $\bar{\sigma}$ acting on all walls. As previously mentioned, two compartments of a strained wall will be under different stresses according to their stiffness difference. The stress acting on wall compartment between cells $\alpha$ and $\beta \in \mathcal{N}_\alpha$, facing cell $\alpha$ is

$$\sigma_{\alpha\beta} = \frac{2\bar{\sigma} E_\alpha}{E_\alpha + E_\beta}. \tag{38}$$

Therefore, all stiffnesses being the same, so are all stresses and, consequently, PIN densities. Since we are not going to account for noise in the analytical treatment we are going to set the auxin expression rate $g(\tau)$ to its average value, $\langle g(\tau)\rangle = 1$. With these assumptions and conditions, $a_\alpha = 1$, $\forall\alpha$ is the trivial steady state of the system and no deformations other than pure inflation are present. This also means that for the specific case of a regular lattice, $l_{\alpha\beta}/L_\alpha$ is simply $1/R$, where $R$ is the number of neighbours. Considering a regular lattice we can also factor out $W = L^*_\alpha/A^*_\alpha = (LL_\alpha)/A_\alpha$ as a lattice geometry dependent constant. For an hexagonal lattice of side $L$, $R = 6$ and $W = 4/\sqrt{3}$.
We can now expand auxin concentration around that state considering small perturbations to auxin $\varepsilon_\alpha$, meaning we transform the system into $a_\alpha = 1 + \varepsilon_\alpha$ and, since $\varepsilon_\alpha$ is small, neglect higher order terms. The time evolution of this perturbation is

$$\frac{da_\alpha}{d\tau} = \frac{d\varepsilon_\alpha}{d\tau} = -\varepsilon_\alpha + \frac{\mathcal{D}W}{R}\sum_{\beta\in\mathcal{N}_\alpha}\left(\varepsilon_\beta - \varepsilon_\alpha\right) + \frac{\mathcal{P}W}{R}\sum_{\beta\in\mathcal{N}_\alpha}\left[p_{\beta\alpha}\frac{1+\varepsilon_\beta}{K+1+\varepsilon_\beta} - p_{\alpha\beta}\frac{1+\varepsilon_\alpha}{K+1+\varepsilon_\alpha}\right], \tag{39}$$

where we have yet to linearize the last term.
Expanding in Taylor series around $\varepsilon_\alpha = 0$ we can rewrite

$$\frac{1+\varepsilon_\alpha}{K+1+\varepsilon_\alpha} = \frac{1}{K+1} + \frac{K}{(K+1)^2}\varepsilon_\alpha + O\left(\varepsilon^2\right). \tag{40}$$

The term $p_{\alpha\beta}$ is slightly less straightforward since it depends not only in the stress applied to the corresponding wall, but also on every other wall of cell $\alpha$. Therefore, the Taylor expansion of $p_{\alpha\beta}$ becomes

$$p_{\alpha\beta} = \left[\frac{f\left(\sigma_{\alpha\beta}\right)}{1+\frac{1}{R}\sum_{\gamma\in\mathcal{N}_\alpha} f\left(\sigma_{\alpha\gamma}\right)}\right|_{\{\varepsilon\}=0} + \left.\frac{\partial p_{\alpha\beta}}{\partial\varepsilon_\alpha}\right|_{\{\varepsilon=0\}}\varepsilon_\alpha + \sum_{\gamma\in\mathcal{N}_\alpha}\left.\frac{\partial p_{\alpha\beta}}{\partial\varepsilon_\gamma}\right|_{\{\varepsilon=0\}}\varepsilon_\gamma + O(\varepsilon^2). \tag{41}$$

The first term is simply

$$p_0 = \frac{f\left(\bar{\sigma}\right)}{1+f\left(\bar{\sigma}\right)}, \tag{42}$$

given that at steady state all stresses are equal to $\bar{\sigma}$. For the second and third terms we need the quantity

$$\frac{\partial p_{\alpha\beta}}{\partial\varepsilon_\gamma} = \frac{\frac{\partial f_{\alpha\beta}}{\partial\varepsilon_\gamma}}{1+\frac{1}{R}\sum_{\kappa\in\mathcal{N}_\alpha} f_{\alpha\kappa}} - \frac{1}{R}\sum_{\kappa\in\mathcal{N}_\alpha}\frac{f_{\alpha\beta}\frac{\partial f_{\alpha\kappa}}{\partial\varepsilon_\gamma}}{\left(1+\frac{1}{R}\sum_{\lambda\in\mathcal{N}_\alpha} f_{\alpha\lambda}\right)^2}, \tag{43}$$

where, since stress will be always positive under these assumptions,

$$\frac{\partial f_{\alpha\beta}}{\partial\varepsilon_\gamma} = n\eta^n\sigma_{\alpha\beta}^{n-1}\frac{\partial\sigma_{\alpha\beta}}{\partial\varepsilon_\gamma}, \tag{44}$$

$$\frac{\partial\sigma_{\alpha\beta}}{\partial\varepsilon_\gamma} = \frac{2\bar{\sigma}}{\left(E_\alpha+E_\beta\right)^2}\left(E_\beta\frac{\partial E_\alpha}{\partial\varepsilon_\gamma} - E_\alpha\frac{\partial E_\beta}{\partial\varepsilon_\gamma}\right), \tag{45}$$

and

$$\frac{\partial E_\alpha}{\partial\varepsilon_\gamma} = E_0 r\delta_{\alpha\gamma}\frac{-2m\left(1+\varepsilon_\gamma\right)^{m-1}}{\left(1+\left(1+\varepsilon_\gamma\right)^m\right)^2}, \tag{46}$$

where $\delta_{\alpha\gamma}$ is the Kronecker delta ($\delta_{\alpha\beta} = 1$ when $\alpha = \beta$, 0 otherwise). Since these derivatives are evaluated in steady state, we have

$$\left.\frac{\partial E_\alpha}{\partial\varepsilon_\gamma}\right|_{\{\varepsilon=0\}} = -\frac{1}{2}E_0 mr\delta_{\alpha\gamma}, \tag{47}$$

,

$$\left.\frac{\partial\sigma_{\alpha\beta}}{\partial\varepsilon_\gamma}\right|_{\{\varepsilon=0\}} = \frac{\bar{\sigma}mr}{4}\left(\delta_{\beta\gamma} - \delta_{\alpha\gamma}\right), \tag{48}$$

and,

$$\left.\frac{\partial f_{\alpha\beta}}{\partial\varepsilon_\gamma}\right|_{\{\varepsilon=0\}} = f\left(\bar{\sigma}\right)\frac{nmr}{4}\left(\delta_{\beta\gamma} - \delta_{\alpha\gamma}\right) = \frac{1}{4}f\left(\bar{\sigma}\right)\mathcal{M}\left(\delta_{\beta\gamma} - \delta_{\alpha\gamma}\right), \tag{49}$$

where $\mathcal{M} = nmr$ contains just parameters related to the feedback between mechanics and PIN. Finally,

$$\left.\frac{\partial p_{\alpha\beta}}{\partial \varepsilon_\gamma}\right|_{\{\varepsilon=0\}} = \frac{1}{4}p_0\mathcal{M}\left[\delta_{\beta\gamma} - \delta_{\alpha\gamma} - \frac{p_0}{R}\sum_{\kappa\in\mathcal{N}_\alpha}\left(\delta_{\kappa\gamma} - \delta_{\alpha\gamma}\right)\right] = \frac{1}{4}p_0\mathcal{M}\left[\delta_{\beta\gamma} - \delta_{\alpha\gamma} + p_0\delta_{\alpha\gamma} - \frac{p_0}{R}\left(1 - \delta_{\alpha\gamma}\right)\right], \tag{50}$$

where we simplified $\sum_{k\in\mathcal{N}_\alpha}\delta_{\kappa\gamma} = 1 - \delta_{\alpha\gamma}$, since it is equal to 1 for $\gamma \in \mathcal{N}_\alpha$ or 0 for $\gamma \notin \mathcal{N}_\alpha$ and we are only interested in the two cases $\gamma = \alpha$ or $\gamma \in \mathcal{N}_\alpha$.
Now, substituting in (41), we get

$$p_{\alpha\beta} = p_0\left[1 + \frac{1}{4}\mathcal{M}\left(\left(\varepsilon_\beta - \varepsilon_\alpha\right) + p_0\varepsilon_\alpha - \frac{p_0}{R}\sum_{\gamma\in\mathcal{N}_\alpha}\varepsilon_\gamma\right)\right] + O(\varepsilon^2). \tag{51}$$

The active transport term is

$$\frac{\mathcal{P}Wp_0}{R\,(K+1)}\left(\frac{K}{K+1} - \frac{\mathcal{M}}{2}\left(1 - \frac{p_0}{2}\right)\right)\sum_{\beta\in\mathcal{N}_\alpha}\left(\varepsilon_\beta - \varepsilon_\alpha\right) - \frac{\mathcal{P}\mathcal{M}Wp_0^2}{4R^2\,(K+1)}\sum_{\beta\in\mathcal{N}_\alpha}\left[\sum_{\kappa\in\mathcal{N}_\beta}\varepsilon_\kappa - \sum_{\kappa\in\mathcal{N}_\alpha}\varepsilon_\kappa\right] + O\left(\varepsilon^2\right). \tag{52}$$

Finally, the linear approximation of the time evolution of the perturbation is

$$\frac{d\varepsilon_\alpha}{d\tau} = -\varepsilon_\alpha + \frac{W}{R}\left[\mathcal{D} + \frac{\mathcal{P}p_0}{K+1}\left(\frac{K}{K+1} - \frac{\mathcal{M}}{2}\left(1 - \frac{p_0}{2}\right)\right)\right]\sum_{\beta\in\mathcal{N}_\alpha}\left(\varepsilon_\beta - \varepsilon_\alpha\right) - \frac{\mathcal{P}\mathcal{M}Wp_0^2}{4R^2\,(K+1)}\sum_{\beta\in\mathcal{N}_\alpha}\left[\sum_{\kappa\in\mathcal{N}_\beta}\varepsilon_\kappa - \sum_{\kappa\in\mathcal{N}_\alpha}\varepsilon_\kappa\right], \tag{53}$$

where

$$W = \frac{L^*}{A^*}, \qquad p_0 = \frac{f\left(\bar{\sigma}\right)}{1 + f\left(\bar{\sigma}\right)}, \qquad \mathcal{M} = nmr. \tag{54}$$

In this form the equation system resembles the class of ODEs studied in appendix A of [2], which we will closely follow. Now we expand $\varepsilon_\alpha$ in a fourier series, exchanging the position of the center of cell $\alpha$, $\mathbf{X}_\alpha$, with wave vectors $\mathbf{k}$,

$$\varepsilon_\mathbf{k} = \frac{1}{2\pi}\sum_\alpha \varepsilon_\alpha e^{-i\mathbf{k}\cdot\mathbf{X}_\alpha}. \tag{55}$$

Let the vectors $\mathbf{e}_p$ denote the vectors from the center of cell $\alpha$ to its $p = 1, 2, \ldots, R$ neighbour. We can define the lattice form factor as

$$S\left(\mathbf{k}\right) = \frac{1}{R}\sum_{p=1}^{R} e^{i\mathbf{k}\cdot\mathbf{e}_p} = \frac{1}{R}\sum_{p=1}^{R} e^{-i\mathbf{k}\cdot\mathbf{e}_p} = \frac{2}{R}\sum_{p=1}^{R/2}\cos\left(\mathbf{k}\cdot\mathbf{e}_p\right), \tag{56}$$

where the second and third equalities follows from considering a regular grid where for each direction $p$ there is another directly opposed to it. This quantity is characteristic of the grid alone and we can rewrite it as

$$S\left(\mathbf{k}\right) = \frac{1}{R}\sum_{\beta\in\mathcal{N}_\alpha} e^{i\mathbf{k}\cdot\left(\mathbf{X}_\beta - \mathbf{X}_\alpha\right)} = \frac{1}{R}\sum_{\beta\in\mathcal{N}_\alpha} e^{-i\mathbf{k}\cdot\left(\mathbf{X}_\beta - \mathbf{X}_\alpha\right)}. \tag{57}$$

The temporal evolution of the quantity $\varepsilon_\mathbf{k}$ is

$$\frac{d\varepsilon_\mathbf{k}}{d\tau} = \frac{1}{2\pi}\sum_\alpha e^{-i\mathbf{k}\cdot\mathbf{X}_\alpha}\frac{d\varepsilon_\alpha}{d\tau}, \tag{58}$$

where we know $d\varepsilon_\alpha/d\tau$. We will treat the three terms in (53) separately. The first term is straightforward,

$$\frac{1}{2\pi}\sum_\alpha e^{-i\mathbf{k}\cdot\mathbf{X}_\alpha}\left(-\varepsilon_\alpha\right) = -\varepsilon_\mathbf{k}. \tag{59}$$

The second is[1]

$$\frac{1}{2\pi}\sum_\alpha e^{-i\mathbf{k}\cdot\mathbf{X}_\alpha}\mathcal{A}\sum_{\beta\in\mathcal{N}_\alpha}\left(\varepsilon_\beta - \varepsilon_\alpha\right) = \frac{\mathcal{A}}{2\pi}\left[\sum_\alpha\sum_{\beta\in\mathcal{N}_\alpha} e^{-i\mathbf{k}\cdot\mathbf{X}_\beta}e^{i\mathbf{k}\cdot\left(\mathbf{X}_\beta - \mathbf{X}_\alpha\right)}\varepsilon_\beta - R\varepsilon_\mathbf{k}\right] = \mathcal{A}R\left(S\left(\mathbf{k}\right) - 1\right)\varepsilon_\mathbf{k}, \tag{60}$$

[1]Where we can write

$$\sum_\alpha\sum_{\beta\in\mathcal{N}_\alpha} e^{-i\mathbf{k}\cdot\mathbf{X}_\beta}e^{i\mathbf{k}\cdot\left(\mathbf{X}_\beta - \mathbf{X}_\alpha\right)}\varepsilon_\beta = \sum_\beta \varepsilon_\beta e^{-i\mathbf{k}\cdot\mathbf{X}_\beta}\sum_{\alpha\in\mathcal{N}_\beta} e^{i\mathbf{k}\cdot\left(\mathbf{X}_\beta - \mathbf{X}_\alpha\right)} = RS\left(\mathbf{k}\right)\varepsilon_\mathbf{k}.$$

This amounts to reordering the terms being summed over on the condition the tissue is infinite.

with $\mathcal{A} = \frac{W}{R}\left[\mathcal{D} + \frac{\mathcal{P}p_0}{K+1}\left(\frac{K}{K+1} - \frac{\mathcal{M}}{2}\left(1 - \frac{p_0}{2}\right)\right)\right]$.
The third one is[2]

$$\frac{1}{2\pi}\sum_{\alpha} e^{-i\mathbf{k}\cdot\mathbf{X}_\alpha}\mathcal{B}\sum_{\beta\in\mathcal{N}_\alpha}\left[\sum_{\kappa\in\mathcal{N}_\beta}\varepsilon_\kappa - \sum_{\kappa\in\mathcal{N}_\alpha}\varepsilon_\kappa\right] = \frac{\mathcal{B}}{2\pi}\left[\sum_{\alpha}\sum_{\beta\in\mathcal{N}_\alpha}\sum_{\kappa\in\mathcal{N}_\beta} e^{-i\mathbf{k}\cdot\mathbf{X}_\kappa}e^{i\mathbf{k}\cdot\left(\mathbf{X}_\kappa-\mathbf{X}_\beta\right)}e^{i\mathbf{k}\cdot\left(\mathbf{X}_\beta-\mathbf{X}_\alpha\right)}\varepsilon_\kappa - R^2 S(\mathbf{k})\right]$$
$$= \mathcal{B}R^2 S(\mathbf{k})\left(S(\mathbf{k}) - 1\right)\varepsilon_{\mathbf{k}}, \quad (61)$$

where $\mathcal{B} = -\frac{\mathcal{P}\mathcal{M}Wp_0^2}{4R^2(K+1)}$. Therefore, adding these three terms together and rearranging we get

$$\frac{d\varepsilon_{\mathbf{k}}}{d\tau} = \left[-1 + W\left(S(\mathbf{k}) - 1\right)\left(\mathcal{D} + \frac{\mathcal{P}p_0}{K+1}\left[\frac{K}{K+1} - \frac{\mathcal{M}}{2}\left(1 + \frac{p_0}{2}\left(S(\mathbf{k}) - 1\right)\right)\right]\right)\right]\varepsilon_{\mathbf{k}}. \quad (62)$$

Therefore, the characteristic equation for a given $\mathbf{k}$ is simply an exponential growth or decay with coefficients

$$\lambda_{\mathbf{k}} = -1 + W\left(S(\mathbf{k}) - 1\right)\left(\mathcal{D} + \frac{\mathcal{P}p_0}{K+1}\left[\frac{K}{K+1} - \frac{\mathcal{M}}{2}\left(1 + \frac{p_0}{2}\left(S(\mathbf{k}) - 1\right)\right)\right]\right), \quad (63)$$

or equivalently, the characteristic equation is the eigenvalue problem $d/d\tau\left(\varepsilon_{\mathbf{k}}\right) = \lambda_{\mathbf{k}}\varepsilon_{\mathbf{k}}$ with eigenvalues $\lambda_{\mathbf{k}}$.
Since only $S$ is a function of $\mathbf{k}$ and all other quantities are parameters, we can find what is the value of $S$ that maximizes $\lambda_{\mathbf{k}}$, i.e. the value of $S$ for the most unstable wave vector $\mathbf{k}^*$. This wave vector is the one which grows faster or decays slower meaning that at long times it will be the one that dominates the system. The value of $S^* = S(\mathbf{k}^*)$ obeys

$$\left.\frac{d\lambda_{\mathbf{k}}}{dS}\right|_{\mathbf{k}=\mathbf{k}^*} = W\left[\mathcal{D} + \frac{\mathcal{P}p_0}{K+1}\left(\frac{K}{K+1} - \frac{\mathcal{M}}{2}\left(1 + p_0\left(S^* - 1\right)\right)\right)\right] = 0, \quad (64)$$

or, since in this case the eigenvalues are degenerate due to lattice symmetry, belonging to the set

$$\Omega = \left\{\mathbf{k}|S(\mathbf{k}) = 1 - \frac{\mathcal{M} - \frac{2K}{K+1} - \frac{2\mathcal{D}(K+1)}{\mathcal{P}p_0}}{\mathcal{M}p_0}\right\}. \quad (65)$$

A condition for patterns to exist is that $|S(\mathbf{k})| \le 1$ (by definition of $S$) , i.e.,

$$\mathcal{M} - \frac{2K}{K+1} - \frac{2\mathcal{D}(K+1)}{\mathcal{P}p_0} \ge 0, \quad (66)$$

since $\mathcal{M}p_0 > 0$.
Another crucial condition is that the wave vector coefficient has to grow in order for the system to adopt the corresponding wave pattern, i.e., when the eigenvalue corresponding to the largest wave vector becomes positive, the fixed point we are expanding on becomes unstable and patterns form. Since we know what $S$ should be for these wave vectors, the condition for pattern formation is

$$\lambda_{\mathbf{k}^*} = \lambda(S^*) = -1 + \frac{\mathcal{P}W}{4(K+1)\mathcal{M}}\left[\mathcal{M} - \frac{2K}{K+1} - \frac{2\mathcal{D}(K+1)}{\mathcal{P}p_0}\right]^2 > 0, \quad (67)$$

or rearranging,

$$\left[\mathcal{M} - \frac{2K}{K+1} - \frac{2\mathcal{D}(K+1)}{\mathcal{P}p_0}\right]^2 > \frac{4(K+1)\mathcal{M}}{\mathcal{P}W} > 0. \quad (68)$$

Thus, both conditions merge into just one,

$$\mathcal{M} - \sqrt{\frac{4(K+1)}{\mathcal{P}W}}\sqrt{\mathcal{M}} - \frac{2K}{K+1} - \frac{2\mathcal{D}(K+1)}{\mathcal{P}p_0} > 0, \quad (69)$$

where we always discard conditions where our parameters are negative. We can now solve for $\sqrt{\mathcal{M}}$ since it is just a second order polynomial. We take the only solution which yields $\sqrt{\mathcal{M}} > 0$ and square the result to obtain

$$\mathcal{M} > \frac{K+1}{\mathcal{P}W}\left[1 + \sqrt{1 + 2W\left(\frac{\mathcal{P}K}{(K+1)^2} + \frac{\mathcal{D}}{p_0}\right)}\right]^2. \quad (70)$$

[2]Where we rewrite,

$$\sum_{\alpha}\sum_{\beta\in\mathcal{N}_\alpha}\sum_{\kappa\in\mathcal{N}_\beta} e^{-i\mathbf{k}\cdot\mathbf{X}_\kappa}e^{i\mathbf{k}\cdot\left(\mathbf{X}_\kappa-\mathbf{X}_\beta\right)}e^{i\mathbf{k}\cdot\left(\mathbf{X}_\beta-\mathbf{X}_\alpha\right)}\varepsilon_\kappa = \sum_{\kappa}\varepsilon_\kappa e^{-i\mathbf{k}\cdot\mathbf{X}_\kappa}\sum_{\beta\in\mathcal{N}_\kappa} e^{i\mathbf{k}\cdot\left(\mathbf{X}_\kappa-\mathbf{X}_\beta\right)}\sum_{\alpha\in\mathcal{N}_\beta} e^{i\mathbf{k}\cdot\left(\mathbf{X}_\beta-\mathbf{X}_\alpha\right)} = R^2 S^2(\mathbf{k})\varepsilon_{\mathbf{k}},$$

which also amounts to reordering terms.
